# Developmental synchrony and extraordinary multiplication rates in pathogenic organisms

**DOI:** 10.1101/2024.02.19.580990

**Authors:** Megan A. Greischar, Lauren M. Childs

## Abstract

The multiplication rates of pathogenic organisms influence disease progression, efficacy of immunity and therapeutics, and potential for within-host evolution. Thus, accurate estimates of multiplication rates are essential for biological understanding. We recently showed that common methods for inferring multiplication rates from malaria infection data substantially overestimate true values (i.e., under simulated scenarios), providing context for extraordinarily large estimates in human malaria parasites. A key unknown is whether this bias arises specifically from malaria parasite biology or represents a broader concern. Here we identify the potential for biased multiplication rate estimates across pathogenic organisms with different developmental biology by generalizing a within-host malaria model. We find that diverse patterns of developmental sampling bias—the change in detectability over developmental age—reliably generate overestimates of the fold change in abundance, obscuring not just true growth rates but potentially even whether populations are expanding or declining. This pattern emerges whenever synchrony, the degree to which development is synchronized across the population of pathogenic organisms comprising an infection, decays with time. Only with simulated increases in synchrony do we find noticeable underestimates of multiplication rates. Obtaining robust estimates of multiplication rates may require accounting for diverse patterns of synchrony in pathogenic organisms.

**Subjects:** computational biology, theoretical biology, ecology, developmental biology

## 1 Introduction

Population growth is often measured by the multiplication rate, the fold change in numbers over a generation, and represents a key predictor of ecological and evolutionary outcomes. Species that can achieve higher multiplication rates on limiting resources are expected to competitively exclude others, a pattern that applies to free-living and pathogenic organisms (e.g., algae, [1], malaria parasites [2]). Variants (e.g., strains, haplotypes) that can sustain higher rates of multiplication over their lifespans will become more common through time, displacing slower-multiplying variants. When that variation is heritable, evolution by natural selection can occur. Multiplication rates also influence the extent to which heritable variation exists, since, for a given mutation rate, populations that multiply faster will accrue more mutations. For pathogenic organisms, preventing populations from acquiring new mutations is a major motivation behind the use of high dose antimicrobial drugs to treat infections, which aims to suppress multiplication rates and thereby reduce the odds of *de novo* resistance mutations, though at the cost of maximizing the competitive advantage of any existing resistant mutants [3, 4]. Distinct from the evolutionary consequences, multiplication rates govern pathogenic organisms’ capacity to harm their hosts [5–7], and preventing or slowing multiplication is hence a target of both artificial interventions (drugs, vaccines) and evolved host defenses. Understanding competitive outcomes, evolutionary potential, pathogenicity, and the health impact of interventions and immune responses requires accurate estimates of multiplication rates.

Estimating the fold change in numbers of pathogenic organisms requires at least two samples from an infection, but by practical necessity, those samples represent only a subset of the total population. Pathogenic organisms residing in tissues are especially challenging to sample, but, even for organisms that are accessible to sampling during their development, the resulting multiplication rates may fail to represent the full population (e.g., malaria parasites [8]). Beyond sampling noise, we recently showed that sampling bias related to the timing of development can generate the appearance of extraordinarily large multiplication rates in human malaria infections with *Plasmodium falciparum* [9]. Two aspects of malaria biology combine to give the appearance of spuriously rapid population growth: First, parasites are difficult to sample in the latter portion of their development, when parasite-occupied red blood cells produce a protein that assists in binding to the walls of capillaries (sequestration), thereby avoiding filtration and removal by the spleen [10]. Second, parasite development is often (though not always) synchronized across the population of parasites inhabiting the red blood cells within a host, and the level of synchrony can change through time (reviewed in [11, 12]). If the level of synchrony were constant through time, then even an inability to sample mature parasites would not impact estimates of multiplication rates, but changing levels of synchrony introduce substantial bias (figure 1). Using model simulations, we showed that decaying synchrony can generate erroneously large multiplication rate estimates, explaining why modern and historical human malaria data are reported to have multiplication rates far in excess of values thought to be biologically plausible [9]. These biased estimates make it difficult to distinguish between even vastly different population growth rates.

**Figure 1:**
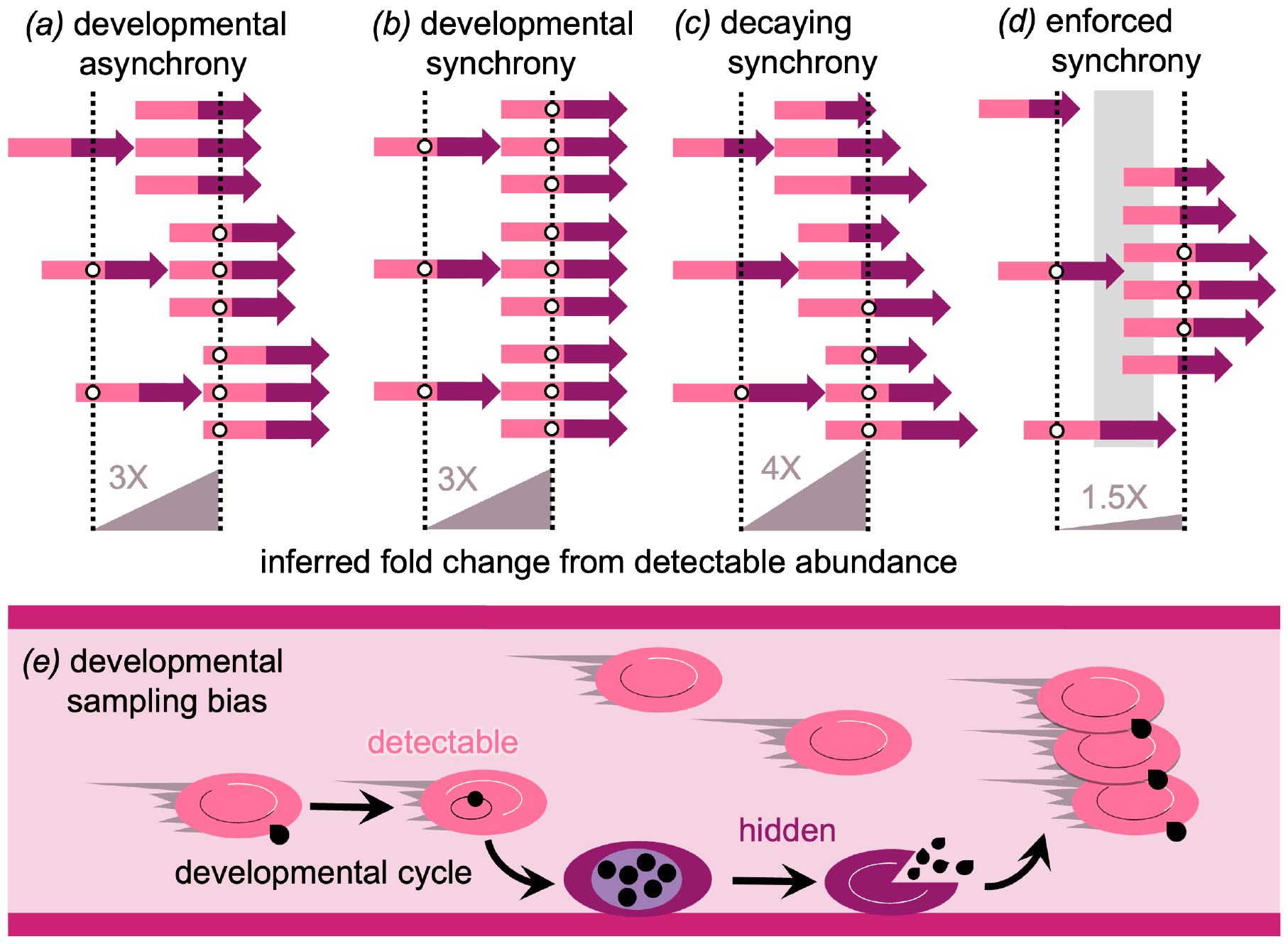
Developmental sampling bias, combined with decaying synchrony, generates overestimates of the fold-change in population abundance. In these examples, developmental progression is indicated with two-toned arrows, individuals are only detectable to sampling in the earlier (light pink) phase of development, and sampling occurs at the mean time required for a developmental cycle. (*a*) When developmental asynchrony is maintained through time, a consistent fraction of the population will be invisible to sampling, yielding accurate estimates of the fold-change in numbers. (*b*) When perfect synchrony is maintained through time, sampling will always catch either all or none of the population. If the population is at a developmentally-detectable age, 3 fold population growth will be correctly inferred. (*c*) When synchrony decays through time, a different fraction of the population will be detectable at successive samples. When the initial sample catches the population with minimal fraction detectable, decaying synchrony ensures that a larger fraction of the population will be detectable in subsequent samples, generating spuriously large estimates of the fold-change in numbers. (*d*) Synchrony can be enforced on an initially asynchronous population, for example if within-host conditions are only amenable to proliferation at discrete intervals (gray area). In that case, sampling can detect a greater fraction of the population initially than at later samples, causing underestimates of the true fold change in abundance. (*e*) The developmental cycle of malaria parasites involves repeated rounds of invasion of and multiplication within the red blood cells of the host. Many malaria parasites exhibit such developmental sampling bias, since red blood cells containing parasites in the early phase of development circulate freely while in the latter phase of development, infected red blood cells adhere to the walls of blood vessels (“sequestration”) in what is thought to be an adaptation to avoid circulation through and removal by the spleen (e.g., [19, 20]).

Given the difficulty of sampling all phases of development equally well *in vivo*, and the variable developmental rhythms observed across pathogenic organisms (reviewed in [13]), these challenges likely arise in diverse systems. Like malaria parasites, *Babesia* parasites multiply in the blood and tend to sequester as development progresses (reviewed in [14]). The blood-borne livestock parasite *Trypanosoma congolense* can also sequester by binding to the blood vessel epithelium for hours at a time [15] and, as with malaria parasites, has been reported to undergo periodic oscillations in detectable abundance in mice [16]. Lymphatic filariasis serves as an extreme example, since the adult worms are sequestered in the lymphatic tissue and only the offspring they produce (microfiliariae) can be sampled in the blood where they circulate, and then only at certain times of day [17]. The timing of sampling is often chosen based on the developmental schedule for the pathogenic organism, as sampling at the same time during the developmental cycle would be expected to yield accurate multiplication rate estimates. For example, sampling of rodent malaria infections is timed so that parasites are early in their development [18]. To understand when and how multiplication rate estimates can be biased by the interplay between sampling and developmental timing, a few key questions must be addressed: (1) what aspects of malaria parasite biology generate grossly inaccurate estimates of population growth rates? (2) in other organisms with different biology and developmental rhythms, to what extent (and in which direction) are estimates of population growth rates likely to be biased? (3) what role does synchrony play in erroneous estimates of population growth rate?

To assess which features of the biology of pathogenic organisms lead to biased estimates of population growth rates, we examine a range of levels of synchrony and variation in the period of detectability. We find that the presence of developmental sampling bias, variation in reliability of sampling as a function of developmental age (i.e., some phases are hidden), is central to the formation of inaccurate assessments of growth rate. However, the extent and direction of the errors in estimated growth rates are determined by the level of synchrony in the population and how that is changing through time. We show that variability in the duration of development, and subsequent minute changes in the age distribution as synchrony decays, make it impossible to sample the population repeatedly at the same point in the developmental cycle. Inaccurate multiplication rate estimates are exacerbated when one of the samples contributing to growth rate estimates occurs at the lowest point of detectability. Next to the situation where the duration of development is invariant, the best case scenario is substantial variation in the duration of development such that synchrony decays rapidly, leading to a more consistent fraction of the population detectable and more accurate multiplication rate estimates. The estimated multiplication rate will depend critically on the initial age distribution of the population of pathogenic organisms, a quantity that is almost never known in advance. These results suggest that understanding variation in developmental synchrony is fundamental to obtaining accurate estimates of multiplication rates.

## 2 Methods

Here, we generalize a model of malaria parasite development to illustrate when and how misleading estimates of population growth rates can arise (figure 1). Malaria is an excellent example as roughly half of the developmental cycle is hidden from sampling and small differences in intrinsic growth rates are magnified during the initial exponential expansion phase of infection. Specifically, we extend a previous model of malaria blood-stage proliferation within the human host [9, 21, 22], to identify the causes and generality of inflated estimates of population growth rates, i.e., the fold-change in abundance from one generation of pathogenic organism to the next. The original model was structured to mimic blood-stage malaria infection, using ordinary differential equations (ODEs) to track the abundance of red blood cells (RBCs) occupied by parasites as they mature through a series of developmental age classes. Development occurs in parallel for parasite-occupied cells that are circulating and available to sampling versus hidden (sequestered). When parameterized from *P. falciparum* data [23], parasite-occupied cells begin to ‘hide’ from sampling (sequester) early in development, with the inflection point occurring at 18.5 hours through the 48 hour window required for parasite development. Under those assumptions, no parasites complete development without sequestering [9, 22], a pattern of developmental sampling bias that—combined with decaying synchrony—leads to spuriously large estimates of the fold increase in the population (figure 1*c*), in excess of 200 when the true (simulated) multiplication rate is only 6-fold [9]. To extend the model beyond malaria biology, we formulate an exemplar baseline pattern for how detectability changes with developmental age and for how synchrony evolves through time. Comparing against this baseline, we simulate with variable relationships between developmental age and detectability as well as diverse patterns of developmental synchrony.

We assume that the mean duration of development is known in advance and sampling occurs as a subset of this development time such that measurements can be used to calculate the fold change at a given time *t* as

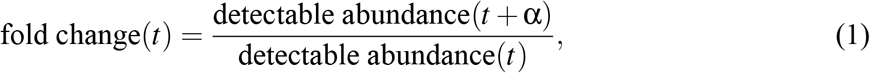

where α represents the mean time required for development. Here, as the original model was motivated by the *P. falciparum* malaria parasite, we assume that development requires two days (α = 2) and that sampling occurs once per day, i.e., twice per developmental cycle. We compare the estimated fold change to the true values, the intrinsic multiplication rate specified in the model (6 fold increase, unless otherwise indicated). We show dynamics for the first three developmental cycles but focus on the fold change from the first and second samples (*t* = 1 and *t* = 2, respectively), since past work suggests that early samples, taken before synchrony has decayed appreciably, are most likely to yield spuriously large estimates of the fold change [9].

### 2.1 Baseline developmental sampling bias

In the baseline scenario, we assume that the pathogenic organisms in the early part of development can be sampled, while those in the latter half of development are hidden, and that the probability of detection ranges from zero to one as the pathogenic organisms develop (as for parasites in malaria infections). We set the inflection point in detectability—where half of the pathogenic organisms are hidden from sampling—to occur halfway through development, rather than at the slightly earlier point as for *P. falciparum* malaria [22]. We assume that the pathogenic organisms transition rapidly from detectable to hidden when they reach the middle of development (slope = 0.2 at the inflection point, which is slightly shallower than that reported for *P. falciparum* in [22]).

Inflated per capita growth rates occur when development is initially synchronized across the population of infected RBCs within the host [9]. We therefore simulate a synchronous start to infections (except where otherwise indicated), with the initial cohort of pathogenic organisms completing development within a nine hour window (that is, the age distribution of the initial cohort is assumed to follow a symmetric beta distribution with both shape parameters set to 100). In addition to the initial level of synchrony, the maintenance of synchrony depends on the variability in developmental duration, with greater variation eroding synchrony more quickly [12]. The compartmental model assumes gamma-distributed duration of development, with the shape parameter of the gamma distribution (*m*) representing the number of compartments (age classes) through which pathogenic organisms must develop before multiplying. The original ODE model tracked parasite development through *m* = 288 age classes, corresponding to a gamma-distributed waiting time around a 48 hour mean developmental duration [22]. Smaller numbers of age classes allow synchrony to decay faster, with the extreme case of only one age compartment corresponding to a standard compartmental model with exponential waiting times. We simulated population dynamics assuming *m* = 200 age compartments as a baseline, allowing for slightly faster decay of synchrony compared with the original malaria-specific model [9, 22].

The severity of overestimated population growth varies considerably with the initial median age of the population, since that determines the fraction of the population that can be detected at regular sampling intervals [9]. If the initial median age were known in advance, it might be possible to adjust sampling times to catch the population when it is mostly detectable, but information on the age structure of populations is rarely available prior to sampling. We therefore simulate a range of initial age distributions with the median ages spanning the full range of development.

### 2.2 Variation in detectability of developmental ages

Assuming the baseline variability in the duration of development, we investigate the impact of changes to the pattern of detectability. First, we assume a slower change in detectability over the course of development, such that the pathogenic organisms transition slowly from detectable to hidden as they progress through development (slope = 0.05 at inflection point). This is in contrast to the baseline case, where most switch from detectable to hidden roughly at the midpoint of development. As a result of the slow change in detectability, some pathogenic organisms are never ‘hidden’ from sampling, while a small fraction enter the hidden state almost immediately.

We then relax the assumption that the pathogenic organisms in the early half of development are most likely to be detected, by reversing the baseline pattern of detectability over developmental ages. The pathogenic organisms instead begin development hidden from sampling and rapidly transition to being detectable at the midpoint of development. None of the pathogenic organisms complete development without becoming detectable. As with the baseline scenario, we simulate populations with a range of initial median ages and estimate the fold increase from the abundance of detectable pathogenic organisms. To assess the impact of population decline, we also simulate baseline conditions and the scenario with late developmental ages detectable using an intrinsic multiplication rate of 1/6.

### 2.3 Diverse patterns of synchrony

Maintaining the baseline pattern of detectability over developmental ages, we investigate the impact of changing synchrony through time. To simulate a more rapid decay in synchrony, we reduce the number of developmental compartments to 50 (compared with *m* = 200 in the baseline case). This doubles the coefficient of variation in the duration of development (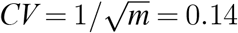 compared with baseline *CV* = 0.07). Greater variation in the duration of development results in faster decay of synchrony, though both this and the baseline scenario exhibit loss of synchrony through time.

To mimic increasing synchrony, we simulate an initially asynchronous population (i.e., a uniform distribution of developmental ages) and enforce synchrony by introducing periodic changes in the intrinsic multiplication rate. Rather than the constant value (6) used in other scenarios, the intrinsic multiplication rate is specified as a sine wave with a period equal to the duration of pathogenic organism development (for simplicity). The sine wave varies from zero to 12 so that the mean intrinsic multiplication rate is 6, the same as other scenarios. Unlike with other scenarios, since we begin with a completely asynchronous population, the initial median age is always the same, and instead vary the phase of the forced sine wave describing the intrinsic multiplication rate. To relate the developmental timing in this case to the initial median age in previous scenarios, we define the forced median age as cycle time (here, 2 days) minus the time of first peak in the intrinsic multiplication rate. For example, if the first peak in the intrinsic multiplication rate occurs at 0.5 days, then pathogenic organisms initially aged 1.5 days will create the largest cohort of daughter organisms. Then the forced median age will tend towards 1.5 days over time, approaching the age distribution of a population beginning the simulation synchronized with an initial median age of 1.5 days.

## 3 Results

### 3.1 Baseline developmental sampling bias

The baseline scenario exhibits substantial overestimation in population growth rates, due to periodic oscillations in the fraction of the population detectable to sampling (figure 2). For a given sampling schedule, the fraction of the population detectable varies considerably depending on the initial median age of the population. Large fluctuations in the fraction detectable become damped over time due to variation in the duration of development and the resulting loss of synchrony (figure 2*a, b*). The calculated fold change in population size is an overestimate when the initial sample (at *t* days) captures a lower fraction detectable than the sample taken one developmental cycle later (*t* + α days, eq. 1). The most severe overestimates of population growth occur when the initial sample represents a small fraction of the population (here corresponding to when the lighter areas line up with sampling times in figure 2*b* at an initial median age of approximately 0.58 days). Due to the decay in synchrony, the abundance sampled one developmental cycle later represents a greater fraction of the population (figure 2*a,b*), akin to the illustration shown in figure 1*c*. Spuriously large estimates also occur when the initial median age is such that the second sample lines up with a low point in the fraction detectable (e.g., when the initial median age is approximately 1.54 days), but the overestimation is less severe because synchrony has already begun to decay, resulting in a greater fraction of the population detectable than previously (figure 2*c*).

**Figure 2:**
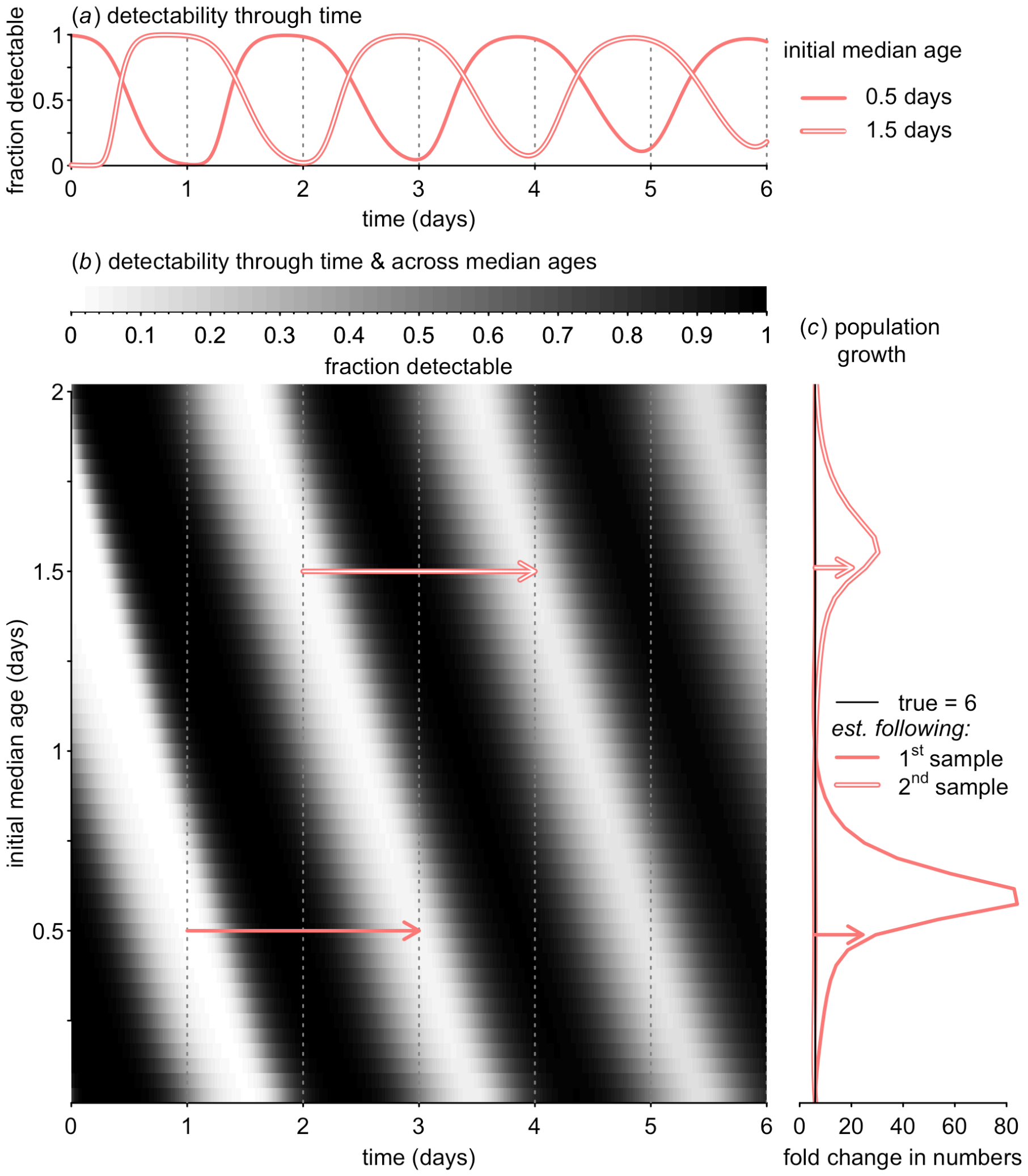
The ability to accurately sample—and infer changes in—abundance varies with the initial median age in the population and the level of developmental synchrony. (*a*) Due to synchrony in the initial cohort, the fraction of the population detectable oscillates through time, shown here for initial median ages of 0.5 and 1.5 days (solid and outlined curve, respectively). Samples were assumed to be collected at day 1, 2, 3, 4, 5 and 6 (dotted vertical lines), and development is assumed to require two days on average. (*b*) The fraction of the population detectable (z axis, color scale) is shown through time (x axis) over the range of possible initial median ages (y axis). Since development requires two days, we assume samples taken two days apart are used to infer the fold change in numbers. When the fraction detectable is low at the initial sample (e.g., at the two trajectories marked with arrows), the inferred fold change in numbers (*c*) represents an overestimate. Note that while modest underestimates of the fold change are possible, the fold change is more commonly (and more severely) overestimated.

With some initial median ages, the fraction detectable is higher at the initial sample than one developmental cycle later. For these initial median ages, population growth is modestly underestimated, with the minimum estimated fold change at approximately 5.3 (true value = 6). While underestimation is possible, overestimation is much more common and severe. For comparison, the worst overestimate in the baseline scenario is an 84-fold expansion. The bias towards over-estimated population growth, while severe, is less extreme than for *P. falciparum*-specific model results, which can exceed 200 when the true value is 6 [9]. The baseline case exhibits less severe bias because we assume detectability declines halfway through development, later than in *P. falciparum* [9, 22, 23], and because we allow synchrony to decay more quickly than previously assumed.

### 3.2 Variation in detectability of developmental ages

We compare the baseline scenario (figure 3*a*) to different scenarios for how detectability changes with developmental age, keeping consistent the variation in the duration of development and hence the rate of decay in synchrony. Overestimation is reduced when changes in detectability are slower and no developmental age is completely hidden (left panels of figure 3*b* versus baseline, figure 3*a*). Since the probability of “being hidden” is lower, it takes longer for the population to reach its minimum detectable fraction compared with the baseline scenario (middle panels of figure 3*b* versus *a*). Thus, the maximal overestimation in the fold-increase occurs at a slightly older initial median age when changes in detectability are slower and incomplete. Both the magnitude of population growth estimation and the initial median age where estimates peak depend on the relationship between development and detectability.

**Figure 3:**
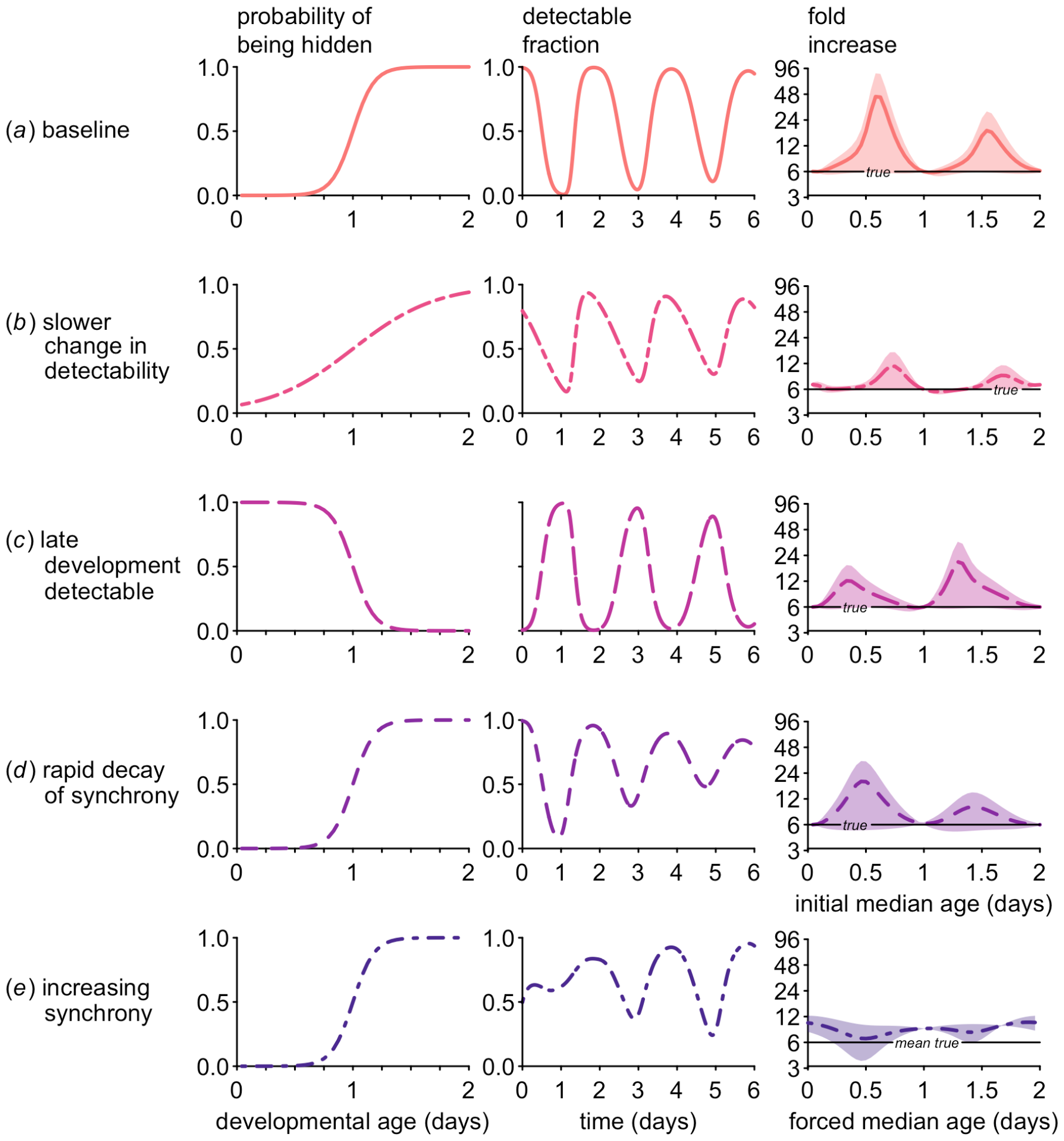
Over a wide range of scenarios, developmental sampling bias can cause substantial overestimates of the fold change of a population. The pattern of developmental sampling bias (i.e., the change in the probability of being hidden from sampling over developmental age, left panels) influences the fraction of the population detectable at sampling (middle panels) and the estimated fold change (right panels). The fraction detectable panels show the change through time for an initial median age of 0.5 days (panels (*a*-*d*) or a forced median age of 0.5 for synchrony enforced on an initially asynchronous population (*e*). In all cases, developmental sampling bias combines with changing synchrony to generate estimates of the fold change in numbers that deviate substantially from the true intrinsic rate of increase specified in the model (6, right panels). For ease of comparison with figure 2, the fold change represents values calculated over two cycles (4 days). (*a*) The baseline case corresponds to the dynamics shown in figure 2*a* (solid line), serving as a comparison for (*b*) when developmental transitions to hidden stages occur more slowly and developmental sampling bias is incomplete; (*c*) when the early (rather than late) phases of development are hidden; (*d*) when synchrony decays rapidly; and finally (*e*) when synchrony increases through time through forcing of multiplication rates. For (*e*), since the multiplication rates vary through time, the black “mean true” line indicates the mean of the sinusoidal multiplication rate function (see Methods for details on forcing and forced median age).

When the early (rather than late) phase of development is hidden (figure 3*c*), changes in the minimal detectable fraction are less rapid because the population is expanding, generating a bias towards earlier developmental ages. As the young developmental ages are hidden, there is a consistently low minimum fraction detectable (middle panel of figure 3*c*). Since the minimal detectable fraction does not change as drastically through time (compared with the baseline scenario) as synchrony decays, the overestimation of population growth is less extreme. The estimated fold-increase from the first sample is maximized when the population begins at an older median age, resulting in most of the population being detectable initially and hence almost none of the population detectable at the first sample (on day 1, see figure 2*b* where, in this reversed scenario, the black areas would indicate a very low fraction of late ages available for sampling). Still, estimated population growth substantially exceeds the true intrinsic values for most initial median ages.

Overestimates of population growth are not isolated to expanding populations, but can also occur when populations are in decline (figure S1), such as after drug treatment. We find that a declining population can appear to be expanding up to 3-fold when late development is detectable (figure S1*b*). Along with the patterns in comparable expanding populations, these results suggest the most extreme overestimates of population growth occur when detectable ages are over-represented in the population, e.g., when late development is detectable in a declining population (figure S1*b*) or when early development is detectable in an expanding population (figure 3*a*). This pattern arises because the detectable fraction of the population is influenced both by the decay of synchrony and by population dynamics (i.e., whether the population is expanding or contracting).

When a population of organisms (pathogenic or otherwise) is growing, individuals in early development will be over-represented, since—by definition—individuals reaching the end of development are more than replacing themselves. Likewise, mature individuals form a disproportionate fraction of declining populations. When synchrony decays through time, the detectable fraction of the population displays damped oscillations such that the minimum detectable fraction increases while the maximum detectable fraction decreases. In an expanding population in which early development can be sampled (figure 3*a*), the minimum detectable fraction increases with time much faster than the maximum detectable fraction declines because early ages are over-represented. The same pattern can be seen for a declining population with late development detectable (figure S1*b*). In both cases, the fold increase can represent a substantial overestimate. In contrast, for an expanding population with late development detectable (figure 3*c*), the minimum detectable fraction increases very slowly, because early ages are over-represented but late ages are all that can be sampled. The minimum detectable fraction also increases slowly in a declining population where only early ages can be sampled (figure S1*a*). In these latter cases, the minimum fraction of the population changes less rapidly through time, reducing (though certainly not eliminating) the problem of over-estimation. Nonetheless, depending on the pattern of detectability across developmental age, and the initial median age of the population, it may not be possible to distinguish population expansion versus decline. To put these results in practical terms, developmental sampling bias may impede ability to estimate not just the magnitude but also the direction of impact of novel therapeutic agents on population growth.

### 3.3 Diverse patterns of synchrony

Retaining the baseline pattern of detectability across developmental age, we find that changes in synchrony impact the direction and magnitude of bias in population growth estimates. Faster decay of synchrony reduces the degree of overestimation in population growth rates (figure 3*d*). In that scenario, the oscillations in the detectable fraction of the population are rapidly damped (middle panel), resulting in better estimates of population growth compared to the baseline scenario (figure 3*a*) where synchrony decays much more slowly. The initial median age at which maximal overestimation of population growth occurs is earlier when synchrony decays faster, a result of the underlying modeling assumption that developmental waiting times are gamma-distributed. While gamma-distributions with a large shape parameter (*m*) are nearly symmetrical, smaller values result in a right-skewed distribution, so that the median lies to the left of the mean (figure S2). As a result, for the more right-skewed distribution, a larger fraction of the population completes development early compared to a more symmetric gamma distribution with a larger shape parameter. Hence, the minimum detectable fraction occurs slightly earlier when synchrony decays rapidly and the most severe overestimation of population growth occurs for a lower initial median age.

When developmental synchrony increases through time as periodicity is forced on an initially asynchronous population (figure 3*e*), calculations of the fold increase can notably underestimate the true value. As with other scenarios, the largest errors in estimation of population growth occur when there are dramatic changes in the minimum fraction of the population detectable. As synchrony is enforced on an asynchronous population, the minimum fraction of the population accessible to sampling decreases with time, leading to underestimated population growth (as illustrated in figure 1*d*, simulated results in figures 3*e* and S3). The most severe underestimates occur when the population is mostly detectable at the first sample and far less detectable one developmental cycle later, which occurs when the forced median age is approximately 0.5 days (figure S3*c*). Even in this case however, overestimates of population growth are still entirely possible, whether the comparison is to the mean intrinsic multiplication rate (as shown in figure 3*e*) or to the true realized multiplication rate, the value calculated as if the entire population were accessible to sampling (left panels, figure S3).

## 4 Discussion

It has long been known that the capacity of a population to expand depends on it age structure [24], including for pathogenic organisms like malaria parasites [19]. Our results show further that the ability to accurately estimate population growth, including whether a population is expanding or declining and how fast, also depends critically on age structure. Across a wide range of biologically plausible scenarios, we find that developmental sampling bias, i.e., variation in the ability to sample individuals at different ages of development, undermines accurate estimation of population growth. Whether early or late development can be sampled, or whether populations are expanding or declining, the bias in multiplication rates is nearly always upwards (figures 3, S1). Multiplication rates can still be accurately estimated *in vitro*, where developmental sampling bias can be minimized or eliminated, but *in vitro* experiments cannot reveal the role of immunity or the impact of vaccines or drugs in limiting the population growth of pathogenic organisms *in vivo*.

A notable exception to multiplication rate overestimates occurs when synchrony increases over time, which can result in marked underestimates of the fold change in numbers. Both increasing and decaying synchrony can occur in the same system. Human malaria infections have been reported to become more synchronous early in infection and also to lose synchrony later in infection [25], though the preponderance of extraordinarily large multiplication rate estimates suggest that the decay of synchrony is very common during the course of malaria infections [9].

The timing of maximal overestimates in population growth rates depends on the initial age distribution of the population and developmental waiting times, both largely uncharted. The number of individuals in each developmental age class (referred to as the initial population vector in matrix population models) remains one of the most poorly communicated quantities across diverse plant and animal species [26] and are rarely known beforehand when sampling infections. Indeed, the developmental sampling bias that makes it difficult to accurately estimate population growth also impedes efforts to characterize the age structure of populations of pathogenic organisms. These knowledge gaps preclude any rules of thumb about the optimal timing of sampling with respect to initial median age to capture population growth.

Better information on developmental synchrony would likely improve estimates of population growth, or at least highlight when they are likely to be erroneously high. Unfortunately, quantifying synchrony is no easy task, even in the well-studied system of malaria infections [12]: common approaches use the percentage of parasites in a particular developmental phase, a metric that will be distorted by both developmental sampling bias and by population growth of the parasites themselves since expanding (declining) populations generate disproportionate percentages of parasites in early (late) development. Taken together with the current results, the presence of developmental sampling bias suggests important benefits to estimating synchrony and population growth jointly rather than independently. An alternative to calculating population growth (eq. 1) is to fit multiplication rates using a dynamic model that includes unsampled compartments (as has been done for HIV data, e.g., [27, 28]). Fitting a model to simultaneously estimate multiplication rates and changing synchrony is likely to require more data than the simpler, but biased calculations. While estimating population growth could require as few as one sample per developmental cycle (eq. 1), quantifying synchrony will require more than one sample per developmental cycle (reviewed in [12]). Therefore, when feasible, it may be advisable to acquire more than one sample per developmental cycle to enable co-estimation of both synchrony and population growth.

Based on estimates of the fold-change from past human malaria and simulated data [9], the largest overestimates of multiplication rates should occur early, before synchrony has had a chance to decay. Accordingly, we find that the first estimated fold increase tends to be larger than the second (figure 2*c*). Since simulated synchrony decays due to the assumed gamma-distributed variation in developmental duration, the minimum detectable fraction increases in successive samples (figure 2*a, b*). Synchrony may arise when pathogenic organisms are under selection to match their developmental timing to host rhythms (reviewed in [13]), including matching circadian rhythms in availability of resources following host meals [29], and timing the completion of incubation periods to coincide with circadian rhythms in host social behavior [30]. Unfortunately, the developmental schedules of pathogenic organisms may have been shaped by evolution in such a way that happens to make estimating multiplication rates incredibly difficult.

These challenges should also arise in free-living organisms, including many insects whose immature stages are inaccessible so that abundance is measured from counts of adults. For example, tea tortrix moth larvae develop within leaves they have webbed together so that population estimates represent counts of adult moths [31]. Similarly, periodical cicadas develop underground before emerging as nymphs, and population estimates rely on counts of these later stages of development [32]. Seasonal temperature variation can increase or reduce variability in the duration of development for insects (including the tea tortrix), altering developmental synchrony [33] and presumably also the ability to accurately estimate multiplication rates. Thus, whenever there is developmental sampling bias, and the level of synchrony is changing through time—both conditions likely to be extraordinarily common across pathogenic and free-living organisms—estimates of the fold change in population abundance will be biased.

## Supporting information

R code for running all analyses, generating all figures

## Acknowledgements

We thank Christina Hernández, Christopher Myers, and Nicole Mideo for useful discussion.

## Author contributions

M.A.G.: conceptualization, investigation, methodology, validation, visualization, writing - original draft; L.M.C.: investigation, methodology, validation, visualization, writing - review & editing. Both authors gave final approval for publication.

## Supplemental figures

**Figure S1:**
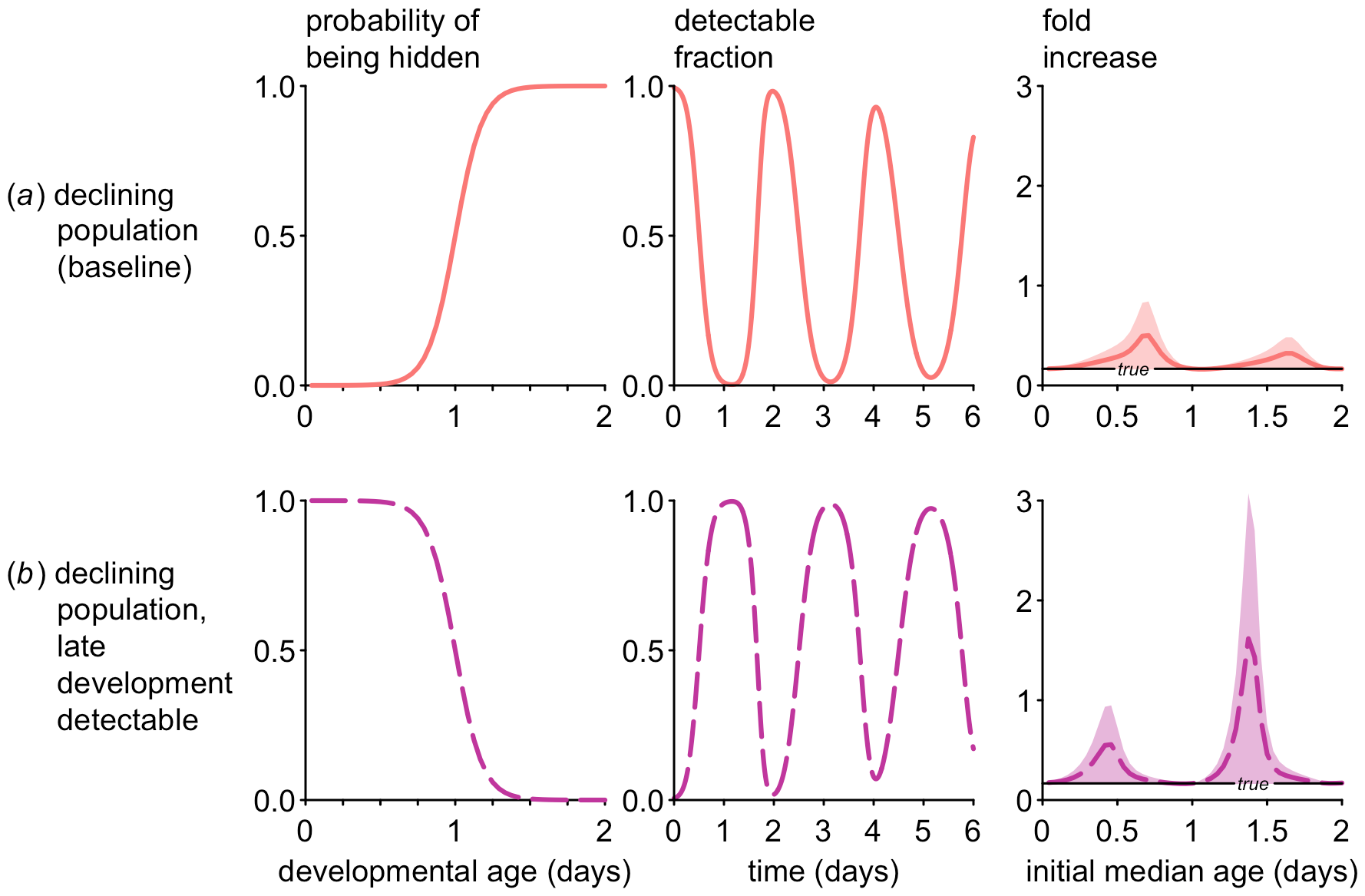
Exaggerated estimates of population growth rates still occur in declining populations. Simulations identical to figure 3*a* and *c* except that the true fold change in abundance is 1/6 rather than 6. Overestimates are relatively minor when late ages are hidden (*a*), and more severe when early ages are hidden (*b*). In the latter case, a population that is actually in decline can appear to be expanding up to three-fold, depending on the initial median age.

**Figure S2:**
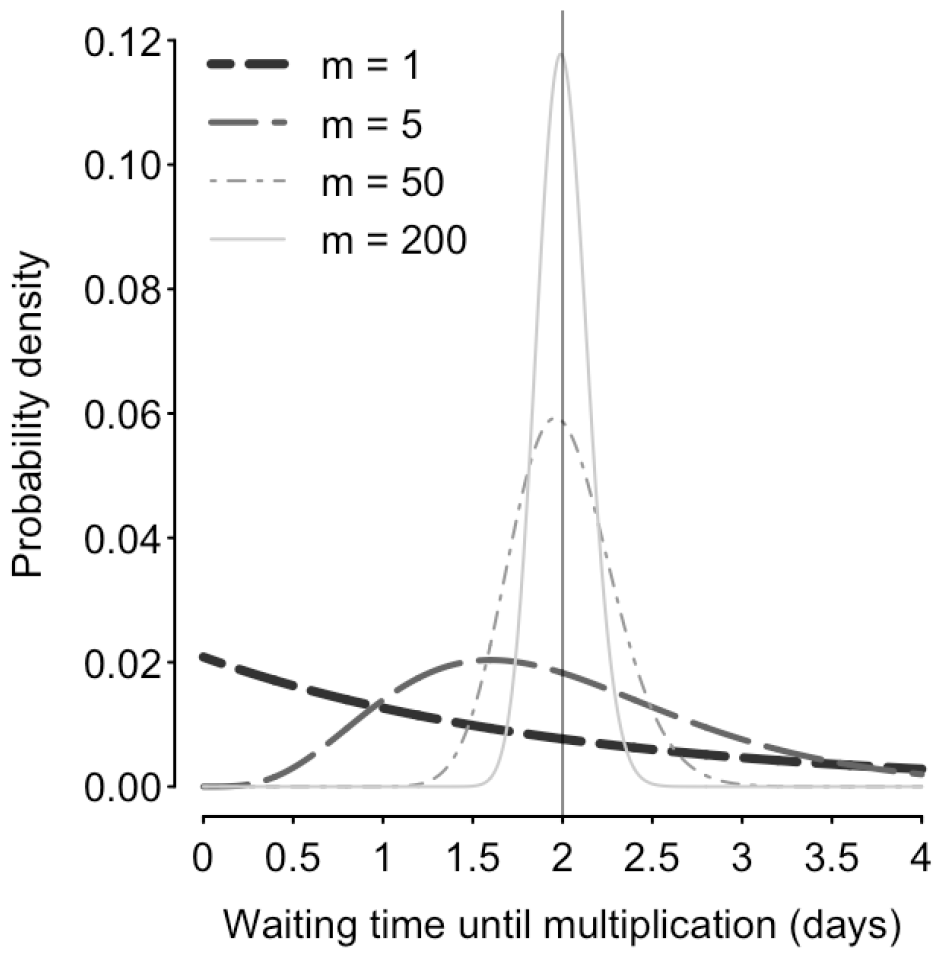
Gamma-distributed waiting times are right-skewed, especially for smaller shape parameters (*m*). The shape parameter represents the number of developmental classes through which pathogenic organisms develop, and the distribution becomes more symmetrical with larger shape parameters (*m*).

**Figure S3:**
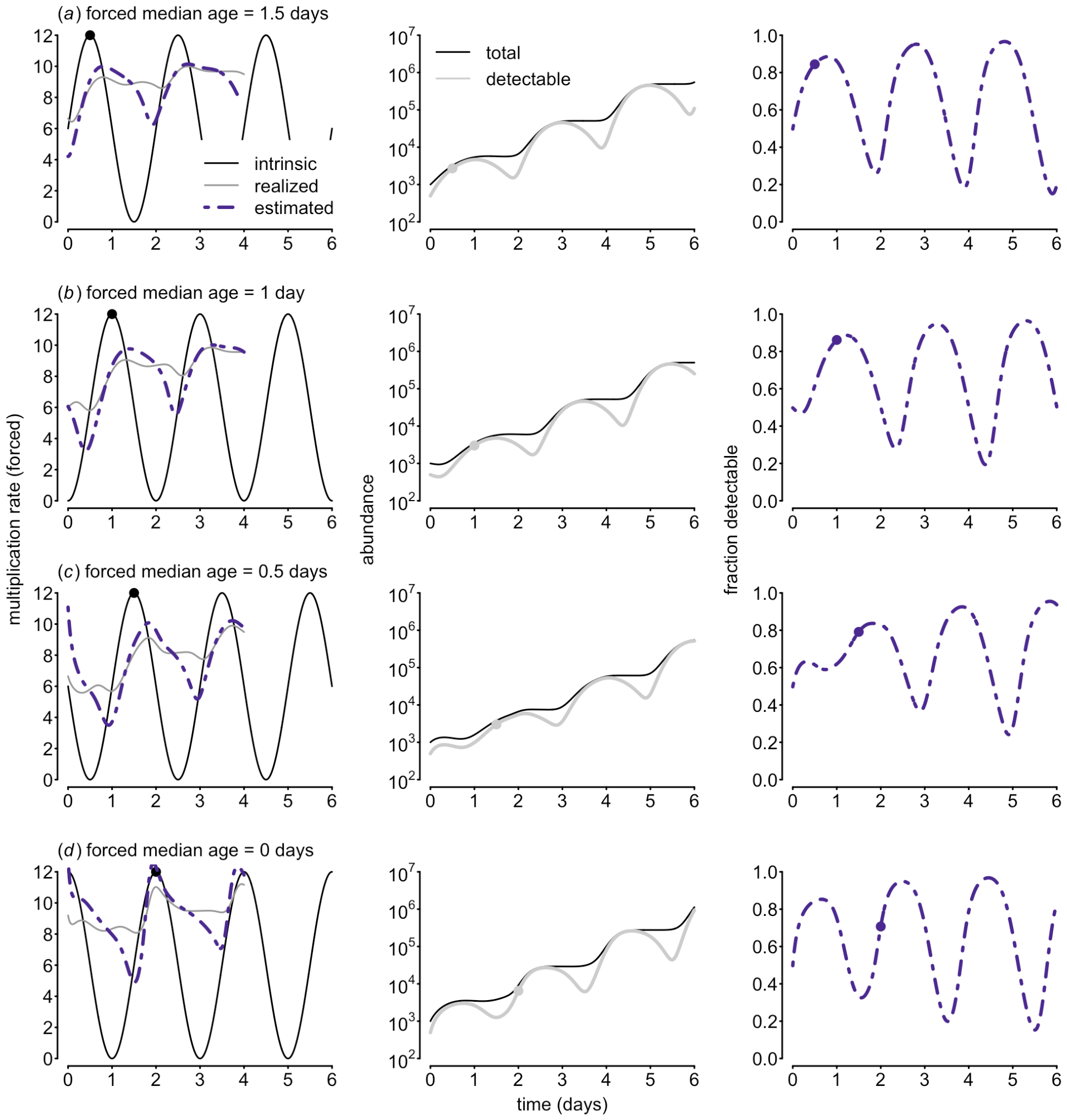
When multiplication rates fluctuate periodically (left panels), it enforces synchrony on an initially asynchronous population (abundance in middle panels and fraction detectable in right panels). Dynamics in (*a*-*d*) simulated assuming different phase shifts in the sinusoidal function used to generate fluctuations in multiplication rates. For reference, the closed dots in each panel correspond to the first peak in multiplication rates, where the forced median age = 2*days* − *time of first peak in multiplication rate*. For example, when the max. multiplication rate occurs at 0.5 days (*a*), the portion of the population aged 1.5 days (what we term the ‘forced median age’) will contribute disproportionately to population growth. Multiplication rates are either intrinsic (the number of daughter cells or virions produced by a mature progenitor), realized (multiplication rates that would be calculated if the entire population could be sampled), or estimated from the detectable abundance.

