## Supplementary material for "Developmental synchrony and extraordinary multiplication rates in pathogenic organisms": R code for running all analyses, generating all figures: SyncBiasedMultRatesCode.R

```
################################
# Code to generate schematic & #
#   calculate PMRs from data.  #
################################

options(warn=1)

library(deSolve)
library(rcartocolor)

# betaOffsetStart returns a vector of the initial iRBC cohort
#   sorted into age compartments according to a Beta distribution
#   where the median age of infection is adjusted from the default
#   (halfway through IED) according to the offset parameter that can
#   take on any value between 0 and 1 (note the function uses the CDF
#   to sort the initial cohort into compartments)
betaOffsetStart = function(compartments, shape, offset, initialI){
  times = seq(0, 1, length.out = round(compartments + 1)) + offset
  correctedTimes = times
  correctedTimes[times > 1] = times[times > 1] - 1
  pbetaValues = pbeta(correctedTimes, shape1 = shape, shape2 = shape)
  correctedpbetaValues = pbetaValues
  correctedpbetaValues[times > 1] = pbetaValues[times > 1] + 1
  iVector = initialI * diff(correctedpbetaValues)
  if(round(sum(iVector)) != round(initialI)){
    warning(paste("betaOffsetStart magnitude error (",
                  round(sum(iVector)) - round(initialI), ", offset = ", 
                  round(offset, digits = 2), ", shape = ", 
                  round(shape, digits = 2), sep = ''), immediate. = T)}
  if(round(length(iVector)) != round(compartments)){
    warning(paste("betaOffsetStart length error, offset = ", 
                  round(offset, digits = 2), ", shape = ", 
                  shape, sep=''), immediate. = T)}
  return(iVector)
}

# function to convert offset value to median parasite age 
# (as fraction through IED)
# multiply by cycle length in hours to get age in hours
offsetToPeak = function(offset){
  res = rep(NA, length(offset))
  for (i in 1:length(offset)){
    res[i] = 1.5 - offset[i]
    if(offset[i]<0.5){res[i] = 0.5 - offset[i]}
  }
  return(res)
}

# function parameterized from Kriek et al. 2003
# for sequestration as a function of hpi
gfx = function(age, p1 = 11.3869/467.6209, 
               p2 = 1,
               p3 = 18.5802, 
               p4 = 0.2242){
  gVal = p1+((p2-p1)/(1+10^(p4*(p3-age))))
  return(gVal)
} # end gfx function

# Archer model modified to enable different number of compartments, non-P. falciparum parameters
modArcher=function(t, y, parms){
  # Archer model with constant PMR, variable number of age compartments
  cycleLength = parms[1] # in hours
  mu = parms[2] # mortality of non-sequestered iRBCs
  museq = parms[3] # mortality of sequestered iRBCs
  R = parms[4] # parasite multiplication rate (constant)
  m = parms[5] # age compartments for intraerythrocytic development (IED)
  min.p = parms[6] # p1
  max.p = parms[7] # p2
  inf.pt = parms[8] # p3
  slope.p = parms[9] # p4
  
  lambdaN = m/cycleLength
  lambdaS = lambdaN
  
  # generate vector of ages through IED:
  ageValues = seq(1/lambdaN,cycleLength,by=1/lambdaN) 
  gValues = gfx(ageValues, p1 = min.p, p2 = max.p, p3 = inf.pt, p4 = slope.p)
  yValues = 1-gValues
  qValues = c(0, 1 - yValues[2:length(yValues)]/yValues[1:(length(yValues)-1)])
  
  N = y[1:m]
  S = y[(m+1):(2*m)]
  
  dN = rep(0, m)
  dS = rep(0, m)
  dN[1] = R*(lambdaN * N[m] + lambdaS * S[m]) - (lambdaN + mu) * N[1]
  dS[1] = 0
  for (i in 2:m){
    dN[i] = (1-qValues[(i-1)]) * lambdaN * N[(i-1)] - (lambdaN + mu) * N[i]
    dS[i] = qValues[(i-1)] * lambdaN * N[(i-1)] + lambdaS * S[(i-1)] - (lambdaS + museq) * S[i]
  } # end for loop through m compartments
  
  res=c(dN,dS)
  list(res)
} # end simplified gamma chain model

forcedPMR = function(t, maxPMR = 6, phase){
  # note that cycle length is in units of hours
  return(maxPMR/2*sin(pi*(t-phase)/24) + maxPMR/2)
  } # end forcedPMR function

# modified Archer model with forced PMR
modArcherForcedPMR=function(t, y, parms){
  # Archer model with constant PMR, variable number of age compartments
  cycleLength = parms[1] # in hours
  mu = parms[2] # mortality of non-sequestered iRBCs
  museq = parms[3] # mortality of sequestered iRBCs
  Rmax = parms[4] # parasite multiplication rate (constant)
  m = parms[5] # age compartments for intraerythrocytic development (IED)
  min.p = parms[6] # p1
  max.p = parms[7] # p2
  inf.pt = parms[8] # p3
  slope.p = parms[9] # p4
  phaseVal = parms[10] # phase of forced sine wave
  
  lambdaN = m/cycleLength
  lambdaS = lambdaN
  
  R = forcedPMR(t, maxPMR = Rmax, phase = phaseVal)
  
  # generate vector of ages through IED:
  ageValues = seq(1/lambdaN,cycleLength,by=1/lambdaN) 
  gValues = gfx(ageValues, p1 = min.p, p2 = max.p, p3 = inf.pt, p4 = slope.p)
  yValues = 1-gValues
  qValues = c(0, 1 - yValues[2:length(yValues)]/yValues[1:(length(yValues)-1)])
  
  N = y[1:m]
  S = y[(m+1):(2*m)]
  
  dN = rep(0, m)
  dS = rep(0, m)
  dN[1] = R*(lambdaN * N[m] + lambdaS * S[m]) - (lambdaN + mu) * N[1]
  dS[1] = 0
  for (i in 2:m){
    dN[i] = (1-qValues[(i-1)]) * lambdaN * N[(i-1)] - (lambdaN + mu) * N[i]
    dS[i] = qValues[(i-1)] * lambdaN * N[(i-1)] + lambdaS * S[(i-1)] - (lambdaS + museq) * S[i]
  } # end for loop through m compartments
  
  res=c(dN,dS)
  list(res)
} # end simplified gamma chain model

# function to calculate timeing of max PMR from sine forcing function phase
peakPMRCalc = function(phases, peakShift = 12){
  peakTimes = phases + peakShift
  if(any(peakTimes > 48)){
    peakTimes[peakTimes > 48] = peakTimes[peakTimes > 48] - 48}
  return(peakTimes)
  } # end function to calculate peak PMR

# function to locate local minima for analysis
getLocalMin = function(data, whichLocal = NA){
  # obtains local minima of time series data
  # and returns the corresponding index
  # these are the times corresponding to places where the derivative 
  # switches from negative (decreasing) to positive (increasing) 
  minIndex = which(diff(sign(diff(data))) == 2) + 1
  res = minIndex
  if(!is.na(whichLocal)){
    res = minIndex[[whichLocal]]
    } # end conditional: return specific local minimum (e.g., the 1st or 2nd)
  # # if first value is a global minimum, include that:
  # if(data[1]==min(data)){res = c(1,minIndex)}
  return(res)
  }

initAgeRes = 1
initOffsets = 0.5-seq(initAgeRes,48,by=initAgeRes)/48
initOffsets[initOffsets<0] = 1+initOffsets[initOffsets<0]
initAges = 48*offsetToPeak(initOffsets)
phaseValues = 0:47

baseline = expand.grid(case = "base", R = 6, m = 200, p1 = 0, p2 = 1, p3 = 24, p4 = 0.2, offset = initOffsets, phase = NA, stringsAsFactors = F)
lowM = expand.grid(case = "lowM", R = 6, m = 50, p1 = 0, p2 = 1, p3 = 24, p4 = 0.2, offset = initOffsets, phase = NA, stringsAsFactors = F)
sensiSlope = expand.grid(case = "shallow", R = 6, m = 200, p1 = 0, p2 = 1, p3 = 24, p4 = 0.05, offset = initOffsets, phase = NA, stringsAsFactors = F)
decl = expand.grid(case = "decline", R = 1/6, m = 200, p1 = 0, p2 = 1, p3 = 24, p4 = 0.2, offset = initOffsets, phase = NA, stringsAsFactors = F)
incSync = expand.grid(case = "incSync", R = 12, m = 200, p1 = 0, p2 = 1, p3 = 24, p4 = 0.2, offset = 0, phase = phaseValues, stringsAsFactors = F)

parmsToTest = rbind(baseline, sensiSlope, lowM, decl, incSync, deparse.level = 0)
parmsToTest$firstCircPMR = NA
parmsToTest$secondCircPMR = NA
parmsToTest$firstSeqPMR = NA
parmsToTest$secondSeqPMR = NA
parmsToTest$firstTruePMR = NA
parmsToTest$secondTruePMR = NA
parmsToTest$maxCircPMR = NA
parmsToTest$minCircPMR = NA
parmsToTest$meanCircPMR = NA
parmsToTest$maxSeqPMR = NA
parmsToTest$minSeqPMR = NA
parmsToTest$meanSeqPMR = NA

pfCycleLength = 48 # hours
timeValues = seq(0, pfCycleLength*3, by = 1/6) # time vector in hours
mu = 0
museq = 0
baseCircAbund = matrix(NA, nrow = length(timeValues), ncol = length(initAges))
baseSeqAbund = matrix(NA, nrow = length(timeValues), ncol = length(initAges))

for(rowVal in 1:length(parmsToTest[,1])){
  print(rowVal)
  R = parmsToTest$R[rowVal]
  m = parmsToTest$m[rowVal]
  p1 = parmsToTest$p1[rowVal] # min expression
  p2 = parmsToTest$p2[rowVal] # max expression
  p3 = parmsToTest$p3[rowVal] # inflection point
  p4 = parmsToTest$p4[rowVal] # slope at inflection point
  offsetValue = parmsToTest$offset[rowVal]
  phaseValue = parmsToTest$phase[rowVal]
  
  ages = seq(pfCycleLength/m,48,by=pfCycleLength/m)
  gs = gfx(ages, p1 = p1, p2 = p2, p3 = p3, p4 = p4)
  ys = 1-gs
  qs = c(0, 1 - ys[2:length(ys)]/ys[1:(length(ys)-1)])
  
  # for modified Archer model
  if(is.na(phaseValue)){
    startI0CircSeq = betaOffsetStart(m,100,offsetValue,1000)
    startValues = c((1-gs)*startI0CircSeq,gs*startI0CircSeq)
    
    extendedParmValues = c(pfCycleLength, mu, museq, R, m, p1, p2, p3, p4)
    
    gamma.output = as.data.frame(lsoda(startValues, times = timeValues,
                                       func = modArcher,
                                       parms = extendedParmValues))
    } # end conditional: use modified Archer model? (no PMR forcing)
  
  if(!is.na(phaseValue)){
    startI0CircSeq = betaOffsetStart(m,1,offsetValue,1000)
    startValues = c((1-gs)*startI0CircSeq,gs*startI0CircSeq)
    
    extendedParmValues = c(pfCycleLength, mu, museq, R, m, 
                           p1, p2, p3, p4, phaseValue)
    
    gamma.output = as.data.frame(lsoda(startValues, times = timeValues,
                                       func = modArcherForcedPMR,
                                       parms = extendedParmValues))
    }
  
  # calculate circulating iRBC abundance
  circI = apply(gamma.output[ , 2:(m + 1)],1,sum)
  # calculate sequestered iRBC abundance
  seqI = apply(gamma.output[ , (m + 2):(m*2 + 1)],1,sum)
  indexPMRCalc = which((gamma.output$time/24)%in%c(1:4))
  
  peakVal = round(48*offsetToPeak(parmsToTest$offset[rowVal]))
  if(parmsToTest$case[rowVal] == "base"){
    baseCircAbund[ , round(48*offsetToPeak(offsetValue))] = circI
    baseSeqAbund[ , round(48*offsetToPeak(offsetValue))] = seqI
    } # end conditional: check case value and record full dynamics?

  if(peakVal == 12 & parmsToTest$case[rowVal] %in% 
     c("base", "hiM", "lowM", "shallow", "decline")){
    fracCirc = circI/(circI+seqI)
    write.table(data.frame(time = timeValues, fracCirc = fracCirc),
                paste(parmsToTest$case[rowVal], peakVal, "FracCirc.txt", sep = ''))
    } # end conditional: record fraction detectable for modified Archer model?
  if(!is.na(phaseValue)){
    if(phaseValue %in% c(0,12,24,36)){
      fracCirc = circI/(circI+seqI)
      totalPara = circI + seqI
      pmrsTotal = totalPara[(indexPMRCalc[2]):length(totalPara)]/totalPara[1:(length(totalPara)-indexPMRCalc[2]+1)]
      pmrsCirc = circI[(indexPMRCalc[2]):length(circI)]/circI[1:(length(circI)-indexPMRCalc[2]+1)]
      length(pmrsCirc) <- length(fracCirc)
      length(pmrsTotal) <- length(fracCirc)
      write.table(data.frame(time = timeValues, 
                             R = sapply(timeValues, forcedPMR, maxPMR = R, phase = phaseValue),
                             circI = circI, seqI = seqI, fracCirc = fracCirc, 
                             pmrsCirc = pmrsCirc, pmrsTotal = pmrsTotal),
                  paste(parmsToTest$case[rowVal], phaseValue, "CircSeqFracCirc.txt", sep = ''))
      } 
    } # end conditional: record fraction detectable for forced PMR model?
  if(peakVal == 36 & parmsToTest$case[rowVal]=="base"){
    fracCirc = circI/(circI+seqI)
    write.table(data.frame(time = timeValues, fracCirc = fracCirc),
                paste(parmsToTest$case[rowVal], peakVal, "FracCirc.txt", sep = ''))
    } # end conditional: record fraction detectable for base dynamics?

  # calculate PMRs assuming early stages can be sampled
  para = circI[indexPMRCalc]
  allPMRs = para[3:length(para)]/para[1:(length(para)-2)]
  parmsToTest$firstCircPMR[rowVal] = allPMRs[1]
  parmsToTest$secondCircPMR[rowVal] = allPMRs[2]
  parmsToTest$maxCircPMR[rowVal] = max(allPMRs)
  parmsToTest$minCircPMR[rowVal] = min(allPMRs)
  parmsToTest$meanCircPMR[rowVal] = mean(allPMRs)
  
  # calculate PMRs assuming late stages can be sampled
  seqPara = seqI[indexPMRCalc]
  allSeqPMRs = seqPara[3:length(seqPara)]/seqPara[1:(length(seqPara)-2)]
  parmsToTest$firstSeqPMR[rowVal] = allSeqPMRs[1]
  parmsToTest$secondSeqPMR[rowVal] = allSeqPMRs[2]
  parmsToTest$maxSeqPMR[rowVal] = max(allSeqPMRs)
  parmsToTest$minSeqPMR[rowVal] = min(allSeqPMRs)
  parmsToTest$meanSeqPMR[rowVal] = mean(allSeqPMRs)
  
  # true PMRs assuming all stages can be sampled
  allPara = circI[indexPMRCalc] + seqI[indexPMRCalc]
  allTruePMRs = allPara[3:length(allPara)]/allPara[1:(length(allPara)-2)]
  parmsToTest$firstTruePMR[rowVal] = allTruePMRs[1]
  parmsToTest$secondTruePMR[rowVal] = allTruePMRs[2]
  }

write.table(parmsToTest, "parmsToTest.txt")
write.table(baseCircAbund, "baseCircAbund.txt")
write.table(baseSeqAbund, "baseSeqAbund.txt")

# read in results
parmRes = read.table("parmsToTest.txt")
circBaseMatrix = read.table("baseCircAbund.txt")
seqBaseMatrix = read.table("baseSeqAbund.txt")

fracCircBase = as.matrix(circBaseMatrix/(circBaseMatrix+seqBaseMatrix))
range(fracCircBase)

# horizontal color bar
color.hbar <- function(lut, min, max=-min, nticks=11, ticks=seq(min, max, len=nticks), labs = ticks, title='', cexVal=1) {
  #scale = (length(lut))/(max-min)
  
  plot(c(min,max), c(0,4), type='n', bty='n', xaxt='n', xlab='', yaxt='n', ylab='', main=title, xaxs = "i", yaxs = "i")
  axis(1, at=ticks, labels=labs, las=1, cex.axis=cexVal, tcl = -0.05, mgp=c(3,0.1,0))
  axis(1, at=ticks[!is.na(labs)], labels=labs[!is.na(labs)], las=1, 
       cex.axis=cexVal, tcl = -0.1, mgp=c(3,0.1,0))
  for (i in 1:(length(lut))) {
    x0 = ticks[i]
    x1 = ticks[(i+1)]
    rect(x0,0,x1,4, col=lut[i], border=NA)
  }
  box()
  }

# vertical color bar
color.vbar <- function(lut, min, max=-min, nticks=11, ticks=seq(min, max, len=nticks), labs = ticks, title='', cexVal=1) {
  #scale = (length(lut))/(max-min)
  
  plot(c(0,4), c(min,max), type='n', bty='n', xaxt='n', xlab='', yaxt='n', ylab='', main=title, xaxs = "i", yaxs = "i")
  axis(2, at=ticks, labels=labs, las=1, cex.axis=cexVal, tcl = -0.1, mgp=c(3,0.2,0))
  for (i in 1:(length(lut))) {
    y0 = ticks[i]
    y1 = ticks[(i+1)]
    rect(0,y0,4,y1, col=lut[i], border=NA)
  }
  box()
}

breakValues = seq(0,1,by=0.02)
bandw = colorRampPalette(c("white","black"))
ptcex = 0.75

baseCol = ((cartocolors[cartocolors$Name=="ag_Sunset",]$n7)[[1]])[5]
slowCol = ((cartocolors[cartocolors$Name=="ag_Sunset",]$n7)[[1]])[4]
lateCol = ((cartocolors[cartocolors$Name=="ag_Sunset",]$n7)[[1]])[3]
lowMCol = ((cartocolors[cartocolors$Name=="ag_Sunset",]$n7)[[1]])[2]
incSyncCol = ((cartocolors[cartocolors$Name=="ag_Sunset",]$n7)[[1]])[1]
declEarlyCol = baseCol#((cartocolors[cartocolors$Name=="SunsetDark",]$n7)[[1]])[2]
declLateCol = lateCol#((cartocolors[cartocolors$Name=="SunsetDark",]$n7)[[1]])[7]

fracBase = read.table("base12FracCirc.txt")
fracLowM = read.table("lowM12FracCirc.txt")
fracShallow = read.table("shallow12FracCirc.txt")
fracDecline = read.table("decline12FracCirc.txt")
fracIncSync0 = read.table("incSync0CircSeqFracCirc.txt")
fracIncSync12 = read.table("incSync12CircSeqFracCirc.txt")
fracIncSync24 = read.table("incSync24CircSeqFracCirc.txt")
fracIncSync36 = read.table("incSync36CircSeqFracCirc.txt")

cexText = 0.8

fracBase2 = read.table("base36FracCirc.txt")

# determine timing of first and second maximum PMR
baseScenario = parmRes[parmRes$case=="base",]
48*offsetToPeak(baseScenario$offset[which.max(baseScenario$firstCircPMR)])/24
48*offsetToPeak(baseScenario$offset[which.max(baseScenario$secondCircPMR)])/24

png("Fig2.png", height = 7, width = 6, units = "in", res = 300)

rightBorder = 0.73
par(fig = c(0,rightBorder,0.8,1), bty = 'n', oma = c(0,1,1,0), mar = c(2.5,1,0.5,0), xpd = NA)
plot(c(0,6), c(0,1), type = 'n', xaxt = 'n', yaxt = 'n',
     xlim = c(0,6*24), ylim = c(0,1), xlab = '', ylab = '', xaxs = 'i', yaxs = 'i')
segments(x0 = 24*c(1:6), y0 = rep(0,6), y1 = rep(1,6), lty = 3, col = 'gray50')
points(fracBase$time[fracBase$time<=6*24], fracBase$fracCirc[fracBase$time<=6*24], 
       type='l', col = baseCol, lwd = 2)
axis(1, tcl = -0.1, mgp = c(3,0.1,0), at = 24*seq(0,6,by=1), 
     labels = seq(0,6,by=1), cex.axis = cexText)
axis(2, las=2, tcl = -0.1, mgp = c(3,0.2,0), at = seq(0,1,by=0.25),
     labels = c(0, "", 0.5, "", 1), cex.axis = cexText)
points(fracBase2$time[fracBase2$time<=6*24], fracBase2$fracCirc[fracBase2$time<=6*24],
       type = 'l', col = baseCol, lwd = 3)
points(fracBase2$time[fracBase2$time<=6*24], fracBase2$fracCirc[fracBase2$time<=6*24],
       type = 'l', col = 'white', lwd = 1)
legend(x = 6.1*24, y = 1, legend = c(expression(phantom("12 hours")), 
                                     expression(phantom("36 hours"))), 
       lty = 1, lwd = c(2,3), col = baseCol, y.intersp = 1.2, bty = 'n', 
       cex = cexText, border = NA,
       title = expression(phantom("initial median age")))
legend(x = 6.1*24, y = 1, legend = c(expression("0.5 days"), expression("1.5 days")), 
       lty = c(1,1), lwd = c(2,1), col = c(baseCol, 'white'), y.intersp = 1.2, bty = 'n', 
       cex = cexText, border = NA,
       title = expression("initial median age"))
mtext(expression("time (days)"), side=1, line = 1.1, cex = cexText)
mtext("fraction detectable", side = 2, line = 1.2, cex = cexText)
mtext(expression("("*italic("a")*")"~"detectability through time"), 
      line = 0, cex = cexText, adj = 0)

par(new = T, fig = c(0,rightBorder,0,0.74), bty = 'n', mar = c(2.5,1,1.8,0), xpd = F)
image(x = timeValues, y = initAges, z = fracCircBase, 
      col = bandw(length(breakValues)-1),
      breaks = breakValues, xaxt='n', yaxt='n', xlab = '', ylab = '', xlim = c(0,24*6))
axis(1, tcl = -0.1, mgp = c(3,0.1,0), at = 24*seq(0,6,by=1), 
     labels = seq(0,6,by=1), cex.axis = cexText)
axis(2, las=2, tcl = -0.1, mgp = c(3,0.2,0), at = 24*seq(0.25,2,by=0.25),
     labels = c("", 0.5, "", 1, "", 1.5, "", 2), cex.axis = cexText)
mtext("time (days)", side = 1, line = 1, cex = cexText)
mtext("initial median age (days)", side = 2, line = 1.2, cex = cexText)
mtext(expression("("*italic("b")*") detectability through time & across median ages"), line=3, side=3, cex = cexText, adj=0)
firstLocal = apply(fracCircBase, 2, getLocalMin, whichLocal = 1)
secondLocal = apply(fracCircBase, 2, getLocalMin, whichLocal = 2)
abline(v=24*c(1:6),lty=3,col = 'gray50')
arrows(x0 = 24, x1 = 48+24, y0 = 12, length = 0.1, col = baseCol, lwd = 2)
arrows(x0 = 48, x1 = 48+48, y0 = 36, length = 0.1, col = baseCol, lwd = 3)
arrows(x0 = 48, x1 = 48+48, y0 = 36, length = 0.1, col = 'white', lwd = 1)

par(new=T, fig=c(0,rightBorder,0.69,0.8), bty='n', mar=c(2,1,1,0))
tickLabels = breakValues;tickLabels[!signif(breakValues,2)%in%signif(seq(0,1,by=0.1),2)]<-NA
color.hbar(bandw(length(breakValues)-1), min = min(breakValues), 
           max=max(breakValues), ticks=breakValues, labs = tickLabels, cexVal = cexText)
mtext("fraction detectable", side = 1, line = 1, cex = cexText)

par(new=T, fig=c(0.75,1,0,0.74), bty='n', mar = c(2.5,0,1.8,0.5))
plot(0,1, type = 'n', xaxt='n', yaxt='n', xlab = "", ylab = "",
     xaxs = 'i', yaxs = 'i', xlim = c(-5,85), ylim = range(initAges))
points(parmRes$firstCircPMR[parmRes$case=="base"], initAges, type = 'l', 
       col = baseCol, lwd = 2)
points(parmRes$secondCircPMR[parmRes$case=="base"], initAges, type = 'l', 
       col = baseCol, lwd = 3)
points(parmRes$secondCircPMR[parmRes$case=="base"], initAges, type = 'l', 
       col = 'white', lwd = 1)
abline(v=6, lwd=1)
axis(1, tcl = -0.1, mgp = c(3,0.1,0), at = seq(0,80,by=20),
     labels = seq(0,80,by=20), cex.axis = cexText)
mtext("fold change in numbers", side = 1, line = 1, cex = cexText)
arrows(x0 = 6, 
       x1 = (parmRes$firstCircPMR[parmRes$case=="base"])[which(48*offsetToPeak(parmRes$offset[parmRes$case=="base"])==12)]-5, 
       y0 = 12, length = 0.1, col = baseCol, lwd = 2)
arrows(x0 = 6, 
       x1 = (parmRes$secondCircPMR[parmRes$case=="base"])[which(48*offsetToPeak(parmRes$offset[parmRes$case=="base"])==36)]-5, 
       y0 = 36, length = 0.1, col = baseCol, lwd = 3)
arrows(x0 = 6, 
       x1 = (parmRes$secondCircPMR[parmRes$case=="base"])[which(48*offsetToPeak(parmRes$offset[parmRes$case=="base"])==36)]-5, 
       y0 = 36, length = 0.1, col = 'white', lwd = 1)
l1=legend(x="right", legend = c(expression(phantom("true")),
                                expression(phantom("following")),
                                expression(phantom("1"^"st"~"sample")), 
                                expression(phantom("2"^"nd"~"sample"))), 
          bty = 'n', col = c('white', 'white', baseCol, baseCol), 
          lwd = c(1,1,2,3), cex = cexText)
l2=legend(x="right", legend = c(expression("true = 6"),
                                expression(phantom("following")),
                                expression("1"^"st"~"sample"), 
                                expression("2"^"nd"~"sample")), 
          bty = 'n', col = c('black','white', baseCol, 'white'), lwd = c(1,1,1,1), 
          cex = cexText)
text(x = l2$rect$left + l2$rect$w/2, y = l2$text$y[2], expression(italic("est. following:")), 
     adj = 0.62, cex = cexText)
mtext(expression("  ("*italic("c")*") population"), side = 3, line = 1.1, cex = cexText, adj=0)
mtext(expression("        growth"), side = 3, line = 0.3, cex = cexText, adj=0)

dev.off()

png("Fig3.png", height = 6.5, width = 6, units = "in", res = 300)

h1 = 0.2
h2 = 0.4
h3 = 0.6
h4 = 0.8
vright = 2/3
vleft = 1/3
panelLabelAdj = -2
alphaTransparency = 100

# first row
par(fig = c(0, vleft, h4, 1), bty='n', oma = c(0,6,2,0), mar = c(2,1,0.5,0.5), xpd=NA)
plot(1,1, type = 'n', xlim = c(0,48), ylim = c(0,1), xaxt='n', yaxt='n', 
     xaxs = 'i', yaxs = 'i', xlab = '', ylab = '')
axis(1, tcl = -0.1, mgp = c(3,0.1,0), at = 24*seq(0,2,by=0.25),
     labels = c(0, "", 0.5, "", 1, "", 1.5, "", 2), cex.axis = cexText)
axis(2, las=2, at = c(0,0.5,1), tcl = -0.1, mgp = c(3,0.2,0), cex.axis = cexText)
text(x = panelLabelAdj*24, y = 0.5, expression("("*italic("a")*")"~"baseline"), cex = cexText, adj = c(0,0.5))
points(initAges, gfx(initAges, 
                     p1 = (parmRes$p1[parmRes$case=="base"])[1], 
                     p2 = (parmRes$p2[parmRes$case=="base"])[1], 
                     p3 = (parmRes$p3[parmRes$case=="base"])[1], 
                     p4 = (parmRes$p4[parmRes$case=="base"])[1]), type = 'l',
       col = baseCol, lwd = 2)
mtext("probability of\nbeing hidden", side = 3, line = 0.2, adj = 0, cex=cexText)

par(new=T, fig = c(vleft,vright,h4,1), bty = 'n')
plot(fracBase$time[fracBase$time<=144]/24, fracBase$fracCirc[fracBase$time<=144], type='l', 
     col = baseCol, lwd = 2, xaxt='n', yaxt='n', xlab = '', ylab = '', xaxs = 'i', yaxs = 'i', 
     ylim = c(0,1))
axis(1, tcl = -0.1, mgp = c(3,0.1,0), cex.axis = cexText)
axis(2, las=2, at = c(0,0.5,1), tcl = -0.1, mgp = c(3,0.2,0), cex.axis = cexText)
mtext("detectable\nfraction", side = 3, line = 0.2, adj = 0, cex=cexText)

par(new=T, fig = c(vright,1,h4,1), bty = 'n')
plot(1, 1, type = 'n',
     xaxs = 'i', yaxs = 'i', xaxt='n', yaxt='n', 
     xlab = "", ylab = "", xlim = c(0,48), ylim = c(0.5,2))
axis(1, tcl = -0.1, mgp = c(3,0.1,0), at = seq(0,48,by=12), labels = seq(0,2,by=0.5), cex.axis = cexText)
axis(2, las=2, tcl = -0.1, mgp = c(3,0.2,0), at = log10(c(3,6,12,24,48,96)), labels = c(3,6,12,24,48,96), cex.axis = cexText)
mtext(expression("fold\nincrease"), side = 3, line = 0.2, adj = 0, cex=cexText)
polygon(c(initAges,rev(initAges)), log10(c(parmRes$minCircPMR[parmRes$case=="base"],
                                           rev(parmRes$maxCircPMR[parmRes$case=="base"]))),
        col = rgb(t(col2rgb(baseCol)), alpha = alphaTransparency, maxColorValue = 255), border = NA)
points(initAges, log10(parmRes$meanCircPMR[parmRes$case=="base"]), type = 'l', 
       col = baseCol, lwd = 2)
points(c(0,0.48*24), log10(c(6,6)), type = 'l')
points(c(0.75*24,48), log10(c(6,6)), type = 'l')
text(x = 0.5*24, y = log10(6), expression(italic("true")), cex = 0.6*cexText, adj = 0)

# second row
par(new=T, fig = c(0, vleft, h3, h4), bty='n')
plot(1,1, type = 'n', xlim = c(0,48), ylim = c(0,1), xaxt='n', yaxt='n', 
     xaxs = 'i', yaxs = 'i', xlab = '', ylab = '')
axis(1, tcl = -0.1, mgp = c(3,0.1,0), at = 24*seq(0,2,by=0.25),
     labels = c(0, "", 0.5, "", 1, "", 1.5, "", 2), cex.axis = cexText)
axis(2, las=2, at = c(0,0.5,1), tcl = -0.1, mgp = c(3,0.2,0), cex.axis = cexText)
points(initAges, gfx(initAges, 
                     p1 = (parmRes$p1[parmRes$case=="shallow"])[1], 
                     p2 = (parmRes$p2[parmRes$case=="shallow"])[1], 
                     p3 = (parmRes$p3[parmRes$case=="shallow"])[1], 
                     p4 = (parmRes$p4[parmRes$case=="shallow"])[1]), type = 'l',
       col = slowCol, lwd = 2, lty = 6)
text(x = panelLabelAdj*24, y = 0.7, expression("("*italic("b")*")"~"slower"), 
     cex = cexText, adj = c(0,0.5))
text(x = panelLabelAdj*24, y = 0.51, expression("      change in"), cex = cexText, adj = c(0,0.5))
text(x = panelLabelAdj*24, y = 0.3, expression("      detectability"), cex = cexText, adj = c(0,0.4))

par(new=T, fig = c(vleft,vright,h3,h4), bty = 'n')
plot(fracBase$time[fracBase$time<=144]/24, fracBase$fracCirc[fracBase$time<=144], type='n', 
     col = baseCol, lwd = 2, xaxt='n', yaxt='n', xlab = '', ylab = '', xaxs = 'i', yaxs = 'i', 
     ylim = c(0,1))
axis(1, tcl = -0.1, mgp = c(3,0.1,0), cex.axis = cexText)
axis(2, las=2, at = c(0,0.5,1), tcl = -0.1, mgp = c(3,0.2,0), cex.axis = cexText)
points(fracShallow$time[fracShallow$time<=144]/24, fracShallow$fracCirc[fracShallow$time<=144], 
       type = 'l', col = slowCol, lwd = 2, lty = 6)

par(new=T, fig = c(vright,1,h3,h4), bty = 'n')
plot(1, 1, type = 'n',
     xaxs = 'i', yaxs = 'i', xaxt='n', yaxt='n', 
     xlab = "", ylab = "", xlim = c(0,48), ylim = c(0.5,2))
axis(1, tcl = -0.1, mgp = c(3,0.1,0), at = seq(0,48,by=12), labels = seq(0,2,by=0.5), cex.axis = cexText)
axis(2, las=2, tcl = -0.1, mgp = c(3,0.2,0), at = log10(c(3,6,12,24,48,96)), labels = c(3,6,12,24,48,96), cex.axis = cexText)
polygon(c(initAges,rev(initAges)), log10(c(parmRes$minCircPMR[parmRes$case=="shallow"],
                                           rev(parmRes$maxCircPMR[parmRes$case=="shallow"]))),
        col = rgb(t(col2rgb(slowCol)), alpha = alphaTransparency, maxColorValue = 255), border = NA)
points(initAges, log10(parmRes$meanCircPMR[parmRes$case=="shallow"]), type = 'l', 
       col = slowCol, lwd = 2, lty = 6)
points(c(0,1.58*24), log10(c(6,6)), type = 'l')
points(c(1.85*24,48), log10(c(6,6)), type = 'l')
text(x = 1.6*24, y = log10(6), expression(italic("true")), cex = 0.6*cexText, adj = 0)

# third row
par(new=T, fig = c(0, vleft, h2, h3), bty='n')
plot(1,1, type = 'n', xlim = c(0,48), ylim = c(0,1), xaxt='n', yaxt='n', 
     xaxs = 'i', yaxs = 'i', xlab = '', ylab = '')
axis(1, tcl = -0.1, mgp = c(3,0.1,0), at = 24*seq(0,2,by=0.25),
     labels = c(0, "", 0.5, "", 1, "", 1.5, "", 2), cex.axis = cexText)
axis(2, las=2, at = c(0,0.5,1), tcl = -0.1, mgp = c(3,0.2,0), cex.axis = cexText)
points(initAges, 1-gfx(initAges, 
                       p1 = (parmRes$p1[parmRes$case=="base"])[1], 
                       p2 = (parmRes$p2[parmRes$case=="base"])[1], 
                       p3 = (parmRes$p3[parmRes$case=="base"])[1], 
                       p4 = (parmRes$p4[parmRes$case=="base"])[1]), type = 'l',
       col = lateCol, lwd = 2, lty = 5)
text(x = panelLabelAdj*24, y = 0.7, expression("("*italic("c")*")"~"late"), 
     cex = cexText, adj = c(0,0.5))
text(x = panelLabelAdj*24, y = 0.5, expression("      development"), cex = cexText, adj = c(0,0.5))
text(x = panelLabelAdj*24, y = 0.3, expression("      detectable"), cex = cexText, adj = c(0,0.4))

par(new=T, fig = c(vleft,vright,h2,h3), bty = 'n')
plot(fracBase$time[fracBase$time<=144]/24, fracBase$fracCirc[fracBase$time<=144], type='n', 
     col = baseCol, lwd = 2, xaxt='n', yaxt='n', xlab = '', ylab = '', xaxs = 'i', yaxs = 'i', 
     ylim = c(0,1))
axis(1, tcl = -0.1, mgp = c(3,0.1,0), cex.axis = cexText)
axis(2, las=2, at = c(0,0.5,1), labels = NA, tcl = -0.1, mgp = c(3,0.2,0), cex.axis = cexText)
points(fracBase$time[fracBase$time<=144]/24, 1-fracBase$fracCirc[fracBase$time<=144], type = 'l', 
       col = lateCol, lwd = 2, lty = 5)

par(new=T, fig = c(vright,1,h2,h3), bty = 'n')
plot(1, 1, type = 'n',
     xaxs = 'i', yaxs = 'i', xaxt='n', yaxt='n', 
     xlab = "", ylab = "", xlim = c(0,48), ylim = c(0.5,2))
axis(1, tcl = -0.1, mgp = c(3,0.1,0), at = seq(0,48,by=12), labels = seq(0,2,by=0.5), cex.axis = cexText)
axis(2, las=2, tcl = -0.1, mgp = c(3,0.2,0), at = log10(c(3,6,12,24,48,96)), labels = c(3,6,12,24,48,96), cex.axis = cexText)
polygon(c(initAges,rev(initAges)), log10(c(parmRes$minSeqPMR[parmRes$case=="base"],
                                           rev(parmRes$maxSeqPMR[parmRes$case=="base"]))),
        col = rgb(t(col2rgb(lateCol)), alpha = alphaTransparency, maxColorValue = 255), border = NA)
points(initAges, log10(parmRes$meanSeqPMR[parmRes$case=="base"]), type = 'l', 
       col = lateCol, lwd = 2, lty = 5)
points(c(0,0.28*24), log10(c(6,6)), type = 'l')
points(c(0.55*24,48), log10(c(6,6)), type = 'l')
text(x = 0.3*24, y = log10(6), expression(italic("true")), cex = 0.6*cexText, adj = 0)

# fourth row
par(new=T, fig = c(0, vleft, h1, h2), bty='n')
plot(1,1, type = 'n', xlim = c(0,48), ylim = c(0,1), xaxt='n', yaxt='n', 
     xaxs = 'i', yaxs = 'i', xlab = '', ylab = '')
axis(1, tcl = -0.1, mgp = c(3,0.1,0), at = 24*seq(0,2,by=0.25),
     labels = c(0, "", 0.5, "", 1, "", 1.5, "", 2), cex.axis = cexText)
axis(2, las=2, at = c(0,0.5,1), tcl = -0.1, mgp = c(3,0.2,0), cex.axis = cexText)
points(initAges, gfx(initAges, 
                     p1 = (parmRes$p1[parmRes$case=="lowM"])[1], 
                     p2 = (parmRes$p2[parmRes$case=="lowM"])[1], 
                     p3 = (parmRes$p3[parmRes$case=="lowM"])[1], 
                     p4 = (parmRes$p4[parmRes$case=="lowM"])[1]), type = 'l',
       col = lowMCol, lwd = 2, lty = 2)
text(x = panelLabelAdj*24, y = 0.6, expression("("*italic("d")*")"~"rapid decay"), 
     cex = cexText, adj = c(0,0.5))
text(x = panelLabelAdj*24, y = 0.4, expression("      of synchrony"), cex = cexText, adj = c(0,0.5))

par(new=T, fig = c(vleft,vright,h1,h2), bty = 'n')
plot(fracBase$time[fracBase$time<=144]/24, fracBase$fracCirc[fracBase$time<=144], type='n', 
     col = baseCol, lwd = 2, xaxt='n', yaxt='n', xlab = '', ylab = '', xaxs = 'i', yaxs = 'i', 
     ylim = c(0,1))
axis(1, tcl = -0.1, mgp = c(3,0.1,0), cex.axis = cexText)
axis(2, las=2, at = c(0,0.5,1), tcl = -0.1, mgp = c(3,0.2,0), cex.axis = cexText)
points(fracLowM$time[fracLowM$time<=144]/24, fracLowM$fracCirc[fracLowM$time<=144], type = 'l', 
       col = lowMCol, lwd = 2, lty = 2)

par(new=T, fig = c(vright,1,h1,h2), bty = 'n')
plot(1, 1, type = 'n',
     xaxs = 'i', yaxs = 'i', xaxt='n', yaxt='n', 
     xlab = "", ylab = "", xlim = c(0,48), ylim = c(0.5,2))
axis(1, tcl = -0.1, mgp = c(3,0.1,0), at = seq(0,48,by=12), labels = seq(0,2,by=0.5), cex.axis = cexText)
axis(2, las=2, tcl = -0.1, mgp = c(3,0.2,0), at = log10(c(3,6,12,24,48,96)), labels = c(3,6,12,24,48,96), cex.axis = cexText)
polygon(c(initAges,rev(initAges)), log10(c(parmRes$minCircPMR[parmRes$case=="lowM"],
                                           rev(parmRes$maxCircPMR[parmRes$case=="lowM"]))),
        col = rgb(t(col2rgb(lowMCol)), alpha = alphaTransparency, maxColorValue = 255), border = NA)
points(initAges, log10(parmRes$meanCircPMR[parmRes$case=="lowM"]), type = 'l', 
       col = lowMCol, lwd = 2, lty = 2)
mtext(expression("initial median age (days)"), side=1, line = 1.1, cex = cexText)
points(c(0,0.28*24), log10(c(6,6)), type = 'l')
points(c(0.55*24,48), log10(c(6,6)), type = 'l')
text(x = 0.3*24, y = log10(6), expression(italic("true")), cex = 0.6*cexText, adj = 0)

# fifth row
par(new=T, fig = c(0, vleft, 0, h1), bty='n')
plot(1,1, type = 'n', xlim = c(0,48), ylim = c(0,1), xaxt='n', yaxt='n', 
     xaxs = 'i', yaxs = 'i', xlab = '', ylab = '')
axis(1, tcl = -0.1, mgp = c(3,0.1,0), at = 24*seq(0,2,by=0.25),
     labels = c(0, "", 0.5, "", 1, "", 1.5, "", 2), cex.axis = cexText)
axis(2, las=2, at = c(0,0.5,1), tcl = -0.1, mgp = c(3,0.2,0), cex.axis = cexText)
points(initAges, gfx(initAges, 
                     p1 = (parmRes$p1[parmRes$case=="incSync"])[1], 
                     p2 = (parmRes$p2[parmRes$case=="incSync"])[1], 
                     p3 = (parmRes$p3[parmRes$case=="incSync"])[1], 
                     p4 = (parmRes$p4[parmRes$case=="incSync"])[1]), type = 'l',
       col = incSyncCol, lwd = 2, lty = 4)
text(x = panelLabelAdj*24, y = 0.6, expression("("*italic("e")*")"~"increasing"), 
     cex = cexText, adj = c(0,0.5))
text(x = panelLabelAdj*24, y = 0.4, expression("      synchrony"), cex = cexText, adj = c(0,0.5))
mtext(expression("developmental age (days)"), side=1, line = 1.1, cex = cexText)

par(new=T, fig = c(vleft,vright,0,h1), bty = 'n')
plot(fracIncSync24$time[fracIncSync24$time<=144]/24, fracIncSync24$fracCirc[fracIncSync24$time<=144], type='l', 
     col = incSyncCol, lwd = 2, lty = 4, xaxt='n', yaxt='n', xlab = '', ylab = '', xaxs = 'i', yaxs = 'i', 
     ylim = c(0,1))
axis(1, tcl = -0.1, mgp = c(3,0.1,0), cex.axis = cexText)
axis(2, las=2, at = c(0,0.5,1), tcl = -0.1, mgp = c(3,0.2,0), cex.axis = cexText)
mtext(expression("time (days)"), side=1, line = 1.1, cex = cexText)

par(new=T, fig = c(vright,1,0,h1), bty = 'n')
plot(1, 1, type = 'n',
     xaxs = 'i', yaxs = 'i', xaxt='n', yaxt='n', 
     xlab = "", ylab = "", xlim = c(0,48), ylim = c(0.5,2))
axis(1, tcl = -0.1, mgp = c(3,0.1,0), at = seq(0,48,by=12), labels = seq(0,2,by=0.5), cex.axis = cexText)
axis(2, las=2, tcl = -0.1, mgp = c(3,0.2,0), at = log10(c(3,6,12,24,48,96)), labels = c(3,6,12,24,48,96), cex.axis = cexText)
mtext(expression("forced median age (days)"), side=1, line = 1.1, cex = cexText)
peakPMRVector = 48-peakPMRCalc(parmRes$phase[parmRes$case=="incSync"])
polygon(c(peakPMRVector[order(peakPMRVector)],rev(peakPMRVector[order(peakPMRVector)])), 
        log10(c((parmRes$minCircPMR[parmRes$case=="incSync"])[order(peakPMRVector)],
                rev((parmRes$maxCircPMR[parmRes$case=="incSync"])[order(peakPMRVector)]))),
        col = rgb(t(col2rgb(incSyncCol)), alpha = alphaTransparency, maxColorValue = 255), border = NA)
points(peakPMRVector[order(peakPMRVector)], 
       log10((parmRes$meanCircPMR[parmRes$case=="incSync"])[order(peakPMRVector)]), 
       type = 'l', col = incSyncCol, lwd = 2, lty = 4)
points(c(0,0.73*24), log10(c(6,6)), type = 'l')
points(c(1.33*24,48), log10(c(6,6)), type = 'l')
text(x = 0.75*24, y = log10(6), expression(italic("mean true")), cex = 0.6*cexText, adj = 0)

dev.off()

# declining population
png("FigS1.png", height = 4, width = 6, units = "in", res = 300)

hmiddle = 0.5
vright = 2/3
vleft = 1/3
panelLabelAdj = -2
alphaTransparency = 100

# first row
par(fig = c(0, vleft, hmiddle, 1), bty='n', oma = c(0,6,2,0), mar = c(2,1,0.5,0.5), xpd=NA)
plot(1,1, type = 'n', xlim = c(0,48), ylim = c(0,1), xaxt='n', yaxt='n', 
     xaxs = 'i', yaxs = 'i', xlab = '', ylab = '')
axis(1, tcl = -0.1, mgp = c(3,0.1,0), at = 24*seq(0,2,by=0.25),
     labels = c(0, "", 0.5, "", 1, "", 1.5, "", 2), cex.axis = cexText)
axis(2, las=2, at = c(0,0.5,1), tcl = -0.1, mgp = c(3,0.2,0), cex.axis = cexText)
text(x = panelLabelAdj*24, y = 0.63, expression("("*italic("a")*")"~"declining"), cex = cexText, adj = c(0,0.5))
text(x = panelLabelAdj*24, y = 0.39, expression("      population\n      (baseline)"), cex = cexText, adj = c(0,0.5))
points(initAges, gfx(initAges, 
                     p1 = (parmRes$p1[parmRes$case=="decline"])[1], 
                     p2 = (parmRes$p2[parmRes$case=="decline"])[1], 
                     p3 = (parmRes$p3[parmRes$case=="decline"])[1], 
                     p4 = (parmRes$p4[parmRes$case=="decline"])[1]), type = 'l',
       col = declEarlyCol, lwd = 2, lty = 1)
mtext("probability of\nbeing hidden", side = 3, line = 0.2, adj = 0, cex=cexText)

par(new=T, fig = c(vleft,vright,hmiddle,1), bty = 'n')
plot(fracDecline$time[fracDecline$time<=144]/24, fracDecline$fracCirc[fracDecline$time<=144], type='l', 
     col = declEarlyCol, lwd = 2, lty = 1, xaxt='n', yaxt='n', xlab = '', ylab = '', xaxs = 'i', yaxs = 'i', 
     ylim = c(0,1))
axis(1, tcl = -0.1, mgp = c(3,0.1,0), cex.axis = cexText)
axis(2, las=2, at = c(0,0.5,1), tcl = -0.1, mgp = c(3,0.2,0), cex.axis = cexText)
mtext("detectable\nfraction", side = 3, line = 0.2, adj = 0, cex=cexText)

par(new=T, fig = c(vright,1,hmiddle,1), bty = 'n')
plot(1, 1, type = 'n',
     xaxs = 'i', yaxs = 'i', xaxt='n', yaxt='n', 
     xlab = "", ylab = "", xlim = c(0,48), ylim = c(0,3))
axis(1, tcl = -0.1, mgp = c(3,0.1,0), at = seq(0,48,by=12), labels = seq(0,2,by=0.5), cex.axis = cexText)
axis(2, las=2, tcl = -0.1, mgp = c(3,0.2,0), at = c(0:3), cex.axis = cexText)
mtext(expression("fold\nincrease"), side = 3, line = 0.2, adj = 0, cex=cexText)
polygon(c(initAges,rev(initAges)), c(parmRes$minCircPMR[parmRes$case=="decline"],
                                     rev(parmRes$maxCircPMR[parmRes$case=="decline"])),
        col = rgb(t(col2rgb(declEarlyCol)), alpha = alphaTransparency, maxColorValue = 255), border = NA)
points(initAges, parmRes$meanCircPMR[parmRes$case=="decline"], type = 'l', 
       col = declEarlyCol, lwd = 2)
points(c(0,0.48*24), rep((parmRes$R[parmRes$case=="decline"])[1],2), type = 'l')
points(c(0.75*24,48), rep((parmRes$R[parmRes$case=="decline"])[1],2), type = 'l')
text(x = 0.5*24, y = (parmRes$R[parmRes$case=="decline"])[1], 
     expression(italic("true")), cex = 0.6*cexText, adj = 0)

# second row
par(new=T,fig = c(0, vleft, 0, hmiddle), bty='n', oma = c(0,6,2,0), mar = c(2,1,0.5,0.5), xpd=NA)
plot(1,1, type = 'n', xlim = c(0,48), ylim = c(0,1), xaxt='n', yaxt='n', 
     xaxs = 'i', yaxs = 'i', xlab = '', ylab = '')
axis(1, tcl = -0.1, mgp = c(3,0.1,0), at = 24*seq(0,2,by=0.25),
     labels = c(0, "", 0.5, "", 1, "", 1.5, "", 2), cex.axis = cexText)
axis(2, las=2, at = c(0,0.5,1), tcl = -0.1, mgp = c(3,0.2,0), cex.axis = cexText)
text(x = panelLabelAdj*24, y = 0.75, expression("("*italic("b")*")"~"declining"), cex = cexText, adj = c(0,0.5))
text(x = panelLabelAdj*24, y = 0.26, 
     expression("      population,\n      late\n      development\n      detectable"), cex = cexText, adj = c(0,0.5))
points(initAges, 1-gfx(initAges, 
                       p1 = (parmRes$p1[parmRes$case=="decline"])[1], 
                       p2 = (parmRes$p2[parmRes$case=="decline"])[1], 
                       p3 = (parmRes$p3[parmRes$case=="decline"])[1], 
                       p4 = (parmRes$p4[parmRes$case=="decline"])[1]), type = 'l',
       col = declLateCol, lwd = 2, lty = 5)
mtext(expression("developmental age (days)"), side=1, line = 1.1, cex = cexText)

par(new=T, fig = c(vleft,vright,0,hmiddle), bty = 'n')
plot(fracDecline$time[fracDecline$time<=144]/24, 1-fracDecline$fracCirc[fracDecline$time<=144], type='l', 
     col = declLateCol, lwd = 2, lty = 5, xaxt='n', yaxt='n', xlab = '', ylab = '', xaxs = 'i', yaxs = 'i', 
     ylim = c(0,1))
axis(1, tcl = -0.1, mgp = c(3,0.1,0), cex.axis = cexText)
axis(2, las=2, at = c(0,0.5,1), tcl = -0.1, mgp = c(3,0.2,0), cex.axis = cexText)
mtext(expression("time (days)"), side=1, line = 1.1, cex = cexText)

par(new=T, fig = c(vright,1,0,hmiddle), bty = 'n')
plot(1, 1, type = 'n',
     xaxs = 'i', yaxs = 'i', xaxt='n', yaxt='n', 
     xlab = "", ylab = "", xlim = c(0,48), ylim = c(0,3))
axis(1, tcl = -0.1, mgp = c(3,0.1,0), at = seq(0,48,by=12), labels = seq(0,2,by=0.5), cex.axis = cexText)
axis(2, las=2, tcl = -0.1, mgp = c(3,0.2,0), at = c(0:3), cex.axis = cexText)
polygon(c(initAges,rev(initAges)), c(parmRes$minSeqPMR[parmRes$case=="decline"],
                                     rev(parmRes$maxSeqPMR[parmRes$case=="decline"])),
        col = rgb(t(col2rgb(declLateCol)), alpha = alphaTransparency, maxColorValue = 255), border = NA)
points(initAges, parmRes$meanSeqPMR[parmRes$case=="decline"], type = 'l', 
       col = declLateCol, lwd = 2, lty = 5)
points(c(0,1.28*24), rep((parmRes$R[parmRes$case=="decline"])[1],2), type = 'l')
points(c(1.55*24,48), rep((parmRes$R[parmRes$case=="decline"])[1],2), type = 'l')
text(x = 1.3*24, y = (parmRes$R[parmRes$case=="decline"])[1], 
     expression(italic("true")), cex = 0.6*cexText, adj = 0)
mtext(expression("initial median age (days)"), side=1, line = 1.1, cex = cexText)

dev.off()

cexText = 0.8

png("FigS2.png", height = 3.25, width = 3.25, units = "in", res = 200)

par(mfrow = c(1,1), bty='n', mar = c(2.5,3,0.5,0.5), oma = c(0,0,0,0))
waitingTimes = seq(0,96,length = 1000)
plot(waitingTimes, dgamma(waitingTimes, shape = 1, scale = 48), type = 'l', 
     xaxt = 'n', yaxt='n', col = 'gray20', ylim = c(0,0.12), lwd = 3, lty = 6)
axis(1, at = 24*seq(0, 4, by = 0.5), labels = seq(0, 4, by = 0.5), 
     mgp = c(3,0.1,0), tcl = -0.1, cex.axis = cexText)
axis(2, las = 2, tcl = -0.1, mgp = c(3,0.2,0), cex.axis = cexText)
points(waitingTimes, dgamma(waitingTimes, shape = 5, scale = 48/5), type = 'l', 
       col = 'gray40', lwd = 2, lty = 5)
points(waitingTimes, dgamma(waitingTimes, shape = 50, scale = 48/50), type = 'l', 
       col = 'gray60', lwd = 1, lty = 4)
points(waitingTimes, dgamma(waitingTimes, shape = 200, scale = 48/200), type = 'l', 
       col = 'gray80', lwd = 1, lty = 1)
abline(v=48,lwd=0.5)
legend(x="topleft", legend = c("m = 1", "m = 5", "m = 50", "m = 200"),
       lty = c(6,5,4,1), lwd = c(3,2,1,1), bty='n', y.intersp = 1.2,
       col = c('gray20','gray40','gray60','gray80'), cex = cexText)
mtext("Waiting time until multiplication (days)", side = 1, line = 1.2, cex = cexText)
mtext("Probability density", side = 2, line = 2, cex = cexText)

dev.off()

cexText = 0.8

# dynamics for enforced (increasing) synchrony
png("FigS3.png", height = 7, width = 6.5, units = "in", res = 300)

par(mfrow = c(4,3), bty = 'n', oma = c(2.5,0,0,0), mar = c(0.5,3,3,0.5))
# phase = 0
plot(fracIncSync0$time, fracIncSync0$R, type = 'l', col = 'black', xaxt = 'n', yaxt='n', xlab = '', ylab = '')
legend(x = "bottomright", legend = c("intrinsic","realized","estimated"), col = c('black','gray60',incSyncCol),
       lty = c(1,1,4), lwd = c(1,1,2), bg = 'white', box.col = 'white')
axis(1, tcl = -0.1, mgp = c(3,0.1,0), at = 24*seq(0,(max(timeValues)/24),by = 1), labels = seq(0,(max(timeValues)/24),by=1))
axis(2, tcl = -0.1, mgp = c(3,0.2,0), las=2)
mtext("multiplication rate (forced)", side = 2, line = -1.5, cex = 0.66, outer = T)
points(fracIncSync0$time[fracIncSync0$time%in%c(peakPMRCalc(0))],
       fracIncSync0$R[fracIncSync0$time%in%c(peakPMRCalc(0))], 
       pch = 21, col = 'black', bg = c('black'))
points(fracIncSync0$time, fracIncSync0$pmrsTotal, type = 'l', col = 'gray60')
points(fracIncSync0$time, fracIncSync0$pmrsCirc, type = 'l', col = incSyncCol, lwd = 2, lty = 4)
mtext(expression("("*italic("a")*")"~"forced median age = 1.5 days"), side = 3, line = 0, cex = 0.66, adj = 0)

plot(fracIncSync0$time, log10(fracIncSync0$circI + fracIncSync0$seqI), type='l', xaxt='n', yaxt='n', xlab = '', ylab = '', ylim = c(2,7))
axis(1, tcl = -0.1, mgp = c(3,0.1,0), at = 24*seq(0,(max(timeValues)/24),by = 1), labels = seq(0,(max(timeValues)/24),by=1))
axis(2, tcl = -0.1, mgp = c(3,0.2,0), at = 2:7, 
     labels = c(expression(10^2),expression(10^3),expression(10^4),expression(10^5),expression(10^6),expression(10^7)),las=2)
mtext("abundance", side = 2, line = -17.5, cex = 0.66, outer = T)
points(fracIncSync0$time, log10(fracIncSync0$circI), type = 'l', col = 'gray80', lwd = 2)
points(fracIncSync0$time[fracIncSync0$time%in%c(peakPMRCalc(0))],
       log10(fracIncSync0$circI[fracIncSync0$time%in%c(peakPMRCalc(0))]), 
       pch = 21, col = 'gray80', bg = c('gray80'))
legend(x = "topleft", legend = c("total","detectable"), col = c('black','gray80'), lty = 1, bty = 'n', y.intersp = 1.2)

plot(fracIncSync0$time, fracIncSync0$fracCirc, type = 'l', col = incSyncCol, 
     xaxt='n', yaxt='n', xlab = '', ylab = '', lty = 4, ylim = c(0,1), lwd = 2)
axis(1, tcl = -0.1, mgp = c(3,0.1,0), at = 24*seq(0,(max(timeValues)/24),by = 1), labels = seq(0,(max(timeValues)/24),by=1))
axis(2, tcl = -0.1, mgp = c(3,0.2,0), las=2)
mtext("fraction detectable", side = 2, line = -34, cex = 0.66, outer = T)
points(fracIncSync0$time[fracIncSync0$time%in%c(peakPMRCalc(0))],
       fracIncSync0$fracCirc[fracIncSync0$time%in%c(peakPMRCalc(0))], 
       pch = 21, col = incSyncCol, bg = incSyncCol)

# phase = 12
plot(fracIncSync12$time, fracIncSync12$R, type = 'l', col = 'black', xaxt = 'n', yaxt='n', xlab = '', ylab = '')
axis(1, tcl = -0.1, mgp = c(3,0.1,0), at = 24*seq(0,(max(timeValues)/24),by = 1), labels = seq(0,(max(timeValues)/24),by=1))
axis(2, tcl = -0.1, mgp = c(3,0.2,0), las=2)
points(fracIncSync12$time[fracIncSync12$time%in%c(peakPMRCalc(12))],
       fracIncSync12$R[fracIncSync12$time%in%c(peakPMRCalc(12))], 
       pch = 21, col = 'black', bg = c('black'))
points(fracIncSync12$time, fracIncSync12$pmrsTotal, type = 'l', col = 'gray60')
points(fracIncSync12$time, fracIncSync12$pmrsCirc, type = 'l', col = incSyncCol, lwd = 2, lty = 4)
mtext(expression("("*italic("b")*")"~"forced median age = 1 day"), side = 3, line = 0, cex = 0.66, adj = 0)

plot(fracIncSync12$time, log10(fracIncSync12$circI + fracIncSync12$seqI), type='l', xaxt='n', yaxt='n', xlab = '', ylab = '', ylim = c(2,7))
axis(1, tcl = -0.1, mgp = c(3,0.1,0), at = 24*seq(0,(max(timeValues)/24),by = 1), labels = seq(0,(max(timeValues)/24),by=1))
axis(2, tcl = -0.1, mgp = c(3,0.2,0), at = 2:7, 
     labels = c(expression(10^2),expression(10^3),expression(10^4),expression(10^5),expression(10^6),expression(10^7)),las=2)
points(fracIncSync12$time, log10(fracIncSync12$circI), type = 'l', col = 'gray80', lwd = 2)
points(fracIncSync12$time[fracIncSync12$time%in%c(peakPMRCalc(12))],
       log10(fracIncSync12$circI[fracIncSync12$time%in%c(peakPMRCalc(12))]), 
       pch = 21, col = 'gray80', bg = c('gray80'))

plot(fracIncSync12$time, fracIncSync12$fracCirc, type = 'l', col = incSyncCol, 
     xaxt='n', yaxt='n', xlab = '', ylab = '', lty = 4, ylim = c(0,1), lwd = 2)
axis(1, tcl = -0.1, mgp = c(3,0.1,0), at = 24*seq(0,(max(timeValues)/24),by = 1), labels = seq(0,(max(timeValues)/24),by=1))
axis(2, tcl = -0.1, mgp = c(3,0.2,0), las=2)
points(fracIncSync12$time[fracIncSync12$time%in%c(peakPMRCalc(12))],
       fracIncSync12$fracCirc[fracIncSync12$time%in%c(peakPMRCalc(12))], 
       pch = 21, col = incSyncCol, bg = incSyncCol)

# phase = 24
plot(fracIncSync24$time, fracIncSync24$R, type = 'l', col = 'black', xaxt = 'n', yaxt='n', xlab = '', ylab = '')
axis(1, tcl = -0.1, mgp = c(3,0.1,0), at = 24*seq(0,(max(timeValues)/24),by = 1), labels = seq(0,(max(timeValues)/24),by=1))
axis(2, tcl = -0.1, mgp = c(3,0.2,0), las=2)
points(fracIncSync24$time[fracIncSync24$time%in%c(peakPMRCalc(24))],
       fracIncSync24$R[fracIncSync24$time%in%c(peakPMRCalc(24))], 
       pch = 21, col = 'black', bg = c('black'))
points(fracIncSync24$time, fracIncSync24$pmrsTotal, type = 'l', col = 'gray60')
points(fracIncSync24$time, fracIncSync24$pmrsCirc, type = 'l', col = incSyncCol, lwd = 2, lty = 4)
mtext(expression("("*italic("c")*")"~"forced median age = 0.5 days"), side = 3, line = 0, cex = 0.66, adj = 0)

plot(fracIncSync24$time, log10(fracIncSync24$circI + fracIncSync24$seqI), type='l', xaxt='n', yaxt='n', xlab = '', ylab = '', ylim = c(2,7))
axis(1, tcl = -0.1, mgp = c(3,0.1,0), at = 24*seq(0,(max(timeValues)/24),by = 1), labels = seq(0,(max(timeValues)/24),by=1))
axis(2, tcl = -0.1, mgp = c(3,0.2,0), at = 2:7, 
     labels = c(expression(10^2),expression(10^3),expression(10^4),expression(10^5),expression(10^6),expression(10^7)),las=2)
points(fracIncSync24$time, log10(fracIncSync24$circI), type = 'l', col = 'gray80', lwd = 2)
points(fracIncSync24$time[fracIncSync24$time%in%c(peakPMRCalc(24))],
       log10(fracIncSync24$circI[fracIncSync24$time%in%c(peakPMRCalc(24))]), 
       pch = 21, col = 'gray80', bg = c('gray80'))

plot(fracIncSync24$time, fracIncSync24$fracCirc, type = 'l', col = incSyncCol, 
     xaxt='n', yaxt='n', xlab = '', ylab = '', lty = 4, ylim = c(0,1), lwd = 2)
axis(1, tcl = -0.1, mgp = c(3,0.1,0), at = 24*seq(0,(max(timeValues)/24),by = 1), labels = seq(0,(max(timeValues)/24),by=1))
axis(2, tcl = -0.1, mgp = c(3,0.2,0), las=2)
points(fracIncSync24$time[fracIncSync24$time%in%c(peakPMRCalc(24))],
       fracIncSync24$fracCirc[fracIncSync24$time%in%c(peakPMRCalc(24))], 
       pch = 21, col = incSyncCol, bg = incSyncCol)

# phase = 36
plot(fracIncSync36$time, fracIncSync36$R, type = 'l', col = 'black', xaxt = 'n', yaxt='n', xlab = '', ylab = '')
axis(1, tcl = -0.1, mgp = c(3,0.1,0), at = 24*seq(0,(max(timeValues)/24),by = 1), labels = seq(0,(max(timeValues)/24),by=1))
axis(2, tcl = -0.1, mgp = c(3,0.2,0), las=2)
points(fracIncSync36$time[fracIncSync36$time%in%c(peakPMRCalc(36))],
       fracIncSync36$R[fracIncSync36$time%in%c(peakPMRCalc(36))], 
       pch = 21, col = 'black', bg = c('black'))
points(fracIncSync36$time, fracIncSync36$pmrsTotal, type = 'l', col = 'gray60')
points(fracIncSync36$time, fracIncSync36$pmrsCirc, type = 'l', col = incSyncCol, lwd = 2, lty = 4)
mtext(expression("("*italic("d")*")"~"forced median age = 0 days"), side = 3, line = 0, cex = 0.66, adj = 0)

plot(fracIncSync36$time, log10(fracIncSync36$circI + fracIncSync36$seqI), type='l', xaxt='n', yaxt='n', xlab = '', ylab = '', ylim = c(2,7))
axis(1, tcl = -0.1, mgp = c(3,0.1,0), at = 24*seq(0,(max(timeValues)/24),by = 1), labels = seq(0,(max(timeValues)/24),by=1))
axis(2, tcl = -0.1, mgp = c(3,0.2,0), at = 2:7, 
     labels = c(expression(10^2),expression(10^3),expression(10^4),expression(10^5),expression(10^6),expression(10^7)),las=2)
mtext("time (days)", side = 1, line = 1.2, cex = 0.66)
points(fracIncSync36$time, log10(fracIncSync36$circI), type = 'l', col = 'gray80', lwd = 2)
points(fracIncSync36$time[fracIncSync36$time%in%c(peakPMRCalc(36))],
       log10(fracIncSync36$circI[fracIncSync36$time%in%c(peakPMRCalc(36))]), 
       pch = 21, col = 'gray80', bg = c('gray80'))

plot(fracIncSync36$time, fracIncSync36$fracCirc, type = 'l', col = incSyncCol, 
     xaxt='n', yaxt='n', xlab = '', ylab = '', lty = 4, ylim = c(0,1), lwd = 2)
axis(1, tcl = -0.1, mgp = c(3,0.1,0), at = 24*seq(0,(max(timeValues)/24),by = 1), labels = seq(0,(max(timeValues)/24),by=1))
axis(2, tcl = -0.1, mgp = c(3,0.2,0), las=2)
points(fracIncSync36$time[fracIncSync36$time%in%c(peakPMRCalc(36))],
       fracIncSync36$fracCirc[fracIncSync36$time%in%c(peakPMRCalc(36))], 
       pch = 21, col = incSyncCol, bg = incSyncCol)

dev.off()
```
